# Mechanical memory of confinement pressure governs expansion size in epithelial monolayers

**DOI:** 10.64898/2026.02.23.707597

**Authors:** Linn Engström, Simon K Schnyder, Johannes K Ahnlide, Valeriia Grudtsyna, Martijn Gloerich, Pontus Nordenfelt, Amin Doostmohammadi, Vinay Swaminathan

## Abstract

Epithelial tissues undergo rapid expansion during development, repair, and morphogenesis, yet how tissue-scale growth is coordinated to re-establish homeostasis remains unclear. Here, we show that large epithelial monolayers confined at a wide range of initial densities and mechanochemical states robustly converge to the same final size and density upon release, despite differences in initial cell size, YAP activity, and cell number dynamics. To investigate the underlying mechanism, we combined quantitative experiments with a mechanochemical agent-based model in which mechanical pressure arising from confinement acts as a tissue-scale signal that modulates intracellular cell-cycle activity over time. Using this framework, we show that transient mechanical relaxation during confinement selectively elevates cell-cycle activity in higher-density tissues at the time of release, accelerating early expansion without disrupting final homeostatic outcomes. Together, these results reveal how epithelial tissues coordinate collective growth and robustly restore homeostasis during expansion.

**Significance:** Mechanisms that re-establish homeostasis and coordinate cell proliferation in expanding epithelial tissues such as during development and tissue regeneration remain poorly understood. Here, we used a large 2D epithelial expansion model and agent-based modelling to demonstrate history-dependent regulation of colony growth, mediated by pressure sensing. We find that, typically, equilibration of pressure during confinement just before the onset of expansion results in all expanding tissues of a given starting size reaching a target density and final size independent of starting cell numbers. By transiently reducing actomyosin contractility during confinement, we selectively accelerate the expansion of high-density tissues. Our findings provide key insights into physical mechanisms that govern organ development and tissue repair.

## Introduction

Maintenance of epithelial homeostasis during tissue expansion in embryonic development and wound healing is essential for normal tissue function and repair. This requires tissues to rapidly adapt to increases in size by spatiotemporally regulating proliferation while maintaining appropriate cell numbers and density (1–5). While cell-scale functions such as division and extrusion in this context are well described, how large epithelial tissues composed of asynchronously cycling cells and diverse morphologies coordinate these behaviours across length scales to continuously maintain homeostasis during expansion remains poorly understood (4, 6–12).

Mechanical forces act as regulators of proliferation in epithelial tissues by activating several downstream mechanotransduction pathways (13–17). Stretching can activate YAP, TAZ, and β-catenin to promote cell-cycle entry and progression, while mechanosensitive channels such as Piezo1 respond to both tensile and compressive forces to modulate cell behaviour (11, 13, 14, 16, 18–22). In particular, mechanical activation of YAP has been shown to regulate cell proliferation during lung branching, renewal of the intestinal epithelium and epidermal tissues establishing YAP as a central mediator of local proliferative responses to changes in mechanical stress associated with local cell density (23–26). However, both mechanical stress and cell density vary spatially across expanding tissues, and it remains unclear how such heterogeneous local cues are integrated to activate signals such as YAP/TAZ and coordinate proliferation across the tissue.

Previous experimental and computational studies have shown that coupling proliferation to a tissue pressure, emerging from the interplay between proliferation and tissue expansion, can account for the maintenance of density homeostasis (27–30). However, these models typically idealize intracellular regulation and rely on simplified forms of mechanosensitivity, such as instantaneous reductions in proliferation rate as a function of density, force, or pressure. Thus, it still remains unclear whether and how cells integrate mechanical history to regulate proliferation, how this regulation responds to perturbations, and which intracellular mechanisms confer robustness to such feedback.

To address these questions, we combined controlled tissue-scale experiments with computational modelling. We grew large two-dimensional epithelial monolayers at defined initial densities under circular confinement using PDMS stencils and quantified both cell-scale and tissue-scale properties during subsequent expansion using high-resolution imaging and analysis pipelines. This enabled systematic comparison of expansion dynamics across a wide range of starting densities and mechanochemical states. Using this experimental framework, we find that epithelial tissues robustly re-establish density homeostasis during expansion despite large differences in initial density, cell size, and YAP activity at the time of confinement release. Across conditions, colonies expand to similar final sizes and densities, indicating the presence of tissue-scale coordination mechanisms that buffer local heterogeneity. To investigate the basis of this coordination, we employed an agent-based model incorporating mechanochemical feedback between mechanical pressure and intracellular regulatory state, providing a framework to explore how mechanical history may influence collective tissue behaviour during expansion.

## Results

### Confined epithelial colonies exhibit two density-dependent YAP regimes prior to expansion

To understand the physical mechanisms that regulate the re-establishment of homeostasis after tissue expansion, we first sought to map the relationship between tissue density and the mechanochemical state of confined epithelial monolayers at the onset of expansion. Since YAP activity has previously been shown to be a key indicator and regulator of proliferative potential and mechanotransduction in epithelial tissues, we used the nuclear-to-cytoplasmic ratio of YAP (YAP N/C) as a quantitative readout of tissue mechanochemical state (9, 11, 13, 14, 19). We seeded E-cadherin-RFP–expressing Madin-Darby Canine Kidney (MDCK) cells within 1.5 mm diameter circular PDMS stencils on fibronectin (FN)-coated glass coverslips (Fig. 1A and Supplementary Fig. S1A).

**Figure 1.**
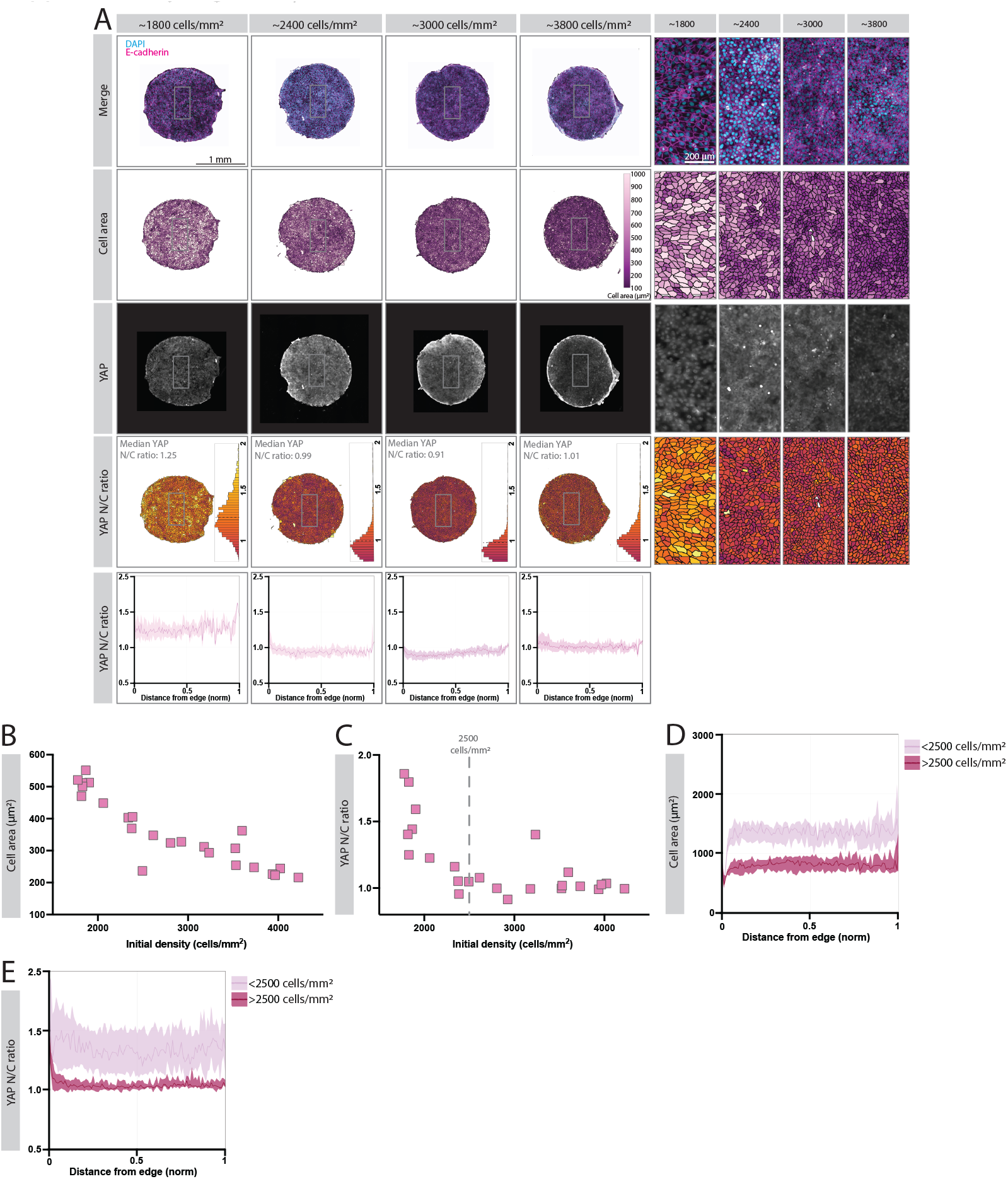
Confined epithelial colonies exhibit two density-dependent YAP regimes prior to expansion. A: Representative images of DAPI/E-cadherin merge, cell area colour maps, YAP channel and YAP N/C colour maps of colonies of varying densities ranging from ∼1800 to ∼3800 cells/mm^2^. Images taken immediately after removal of stencil, color-coded for cell area and YAP N/C ratio. Representative histograms display the population spread for YAP N/C ratios and colour scale from 0.8-2. The dashed lines indicate the median YAP N/C ratio for the colony. Radial spatial plots displaying median YAP N/C ratio of example colonies. Thicker line represents median YAP across colony, with shaded error bars representing interquartile range (IQR) of these. X-axis represents the distance from edge to centre, that has been normalized to arbitrary values. Scale bar whole colony images, 1 mm. Scale bar zoom ins 200 µm. B: Cell area in µm^2^ as a factor of initial density at 0h across the density range ∼1700 cells/mm^2^ to ∼4500 cells/mm^2^ (n = 24 colonies from at least 8 individual experiments). C: YAP N/C ratio as a factor of initial density at 0h across the density range ∼1700 cells/mm^2^ to ∼4500 cells/mm^2^. (n = 24 colonies from at least 8 individual experiments). Dashed line indicates cut off from the YAP-dependent density range (∼1700-2500 cells/mm^2^) to the YAP-independent density range (∼2500-4500 cells/mm^2^) D: Radial spatial plots displaying median cell area of colonies, divided into colonies with a density <2500 cells/mm^2^ and >2500 cells/mm^2^. Thicker line represents median cell area across colony, with shaded error bars representing interquartile range (IQR) of these. X-axis represents the distance from edge to centre, that has been normalized to arbitrary values E: Radial spatial plots displaying median YAP N/C ratio of colonies, divided into colonies with a density <2500 cells/mm^2^ and >2500 cells/mm^2^. Thicker line represents median YAP ratio across colony, with shaded error bars representing interquartile range (IQR) of these. X-axis represents the distance from edge to centre, that has been normalized to arbitrary values

Cells were seeded at varying numbers which led to confined monolayers with densities ranging from ∼1700 to ∼4200 cells/mm^2^ within the stencils. Upon formation of a complete monolayer, stencils were removed, and the colonies were immediately fixed, stained for YAP, F-actin and DAPI, and imaged using high-resolution confocal microscopy. We developed in-house analysis pipelines to generate stitched, single cell resolved maps of the entire colony, enabling precise quantification of cell number, cell area, and YAP N/C for all cells within the colony (Fig. 1A and Supplementary Fig. S1A).

As expected, cell area heat maps showed clear differences across the range of initial colony densities, with a marked reduction in overall cell size in high density colonies compared to lower ones (Fig. 1A). Quantification of median cell area for each colony confirmed a monotonically decreasing trend across the density range (Fig. 1B). In contrast, heat maps of YAP N/C ratio revealed a non-linear dependence on colony density (Fig 1A, Fig 1B). Quantification of median YAP N/C as a function of colony density identified two distinct regimes; in colonies with densities <∼2500 cells/mm^2^, median YAP N/C started high and rapidly decreased with increasing density; while in colonies >∼2500 cells/mm^2^, YAP N/C remained low and was insensitive to further increases in colony density (Fig. 1C and Supplementary Fig. S1B, S1C).

We next asked whether this density-dependent behaviour of cell area and YAP N/C ratio reflected uniform regulation across cells within a colony or instead arose from spatial or cell-to-cell heterogeneity within each density regime. To address this, we analysed YAP N/C distributions and performed spatial analysis of both cell area and YAP N/C within individual colonies. Histograms of YAP N/C in lower density colonies <∼2500 cells/mm^2^ were broad, with a substantial fraction of cells exhibiting high YAP N/C values, whereas distributions in higher density colonies >2500 cells/mm^2^ were significantly narrower and centred around low median YAP N/C values. Spatial analysis further revealed that in low-density colonies, YAP N/C and cell area were uniformly elevated across the colony with no gradient toward the centre, while high-density colonies exhibited a localized increase in YAP N/C restricted to the immediate colony edge, with uniformly low YAP N/C and reduced cell area throughout the colony interior (Fig. 1D, 1E).

### Tissue-wide cell number regulation couples epithelial expansion size to re-establishment of homeostasis

Building on the nonlinear density dependence of YAP activity and the emergence of distinct mechanochemical states prior to expansion, we next investigated how these states influence epithelial expansion after confinement release. To do so, we now allowed the monolayers confined across the density range to expand for 24 or 48 hours prior to fixing, staining and further analysis (Fig. 2A, Supplementary Fig. 2SA).

**Figure 2.**
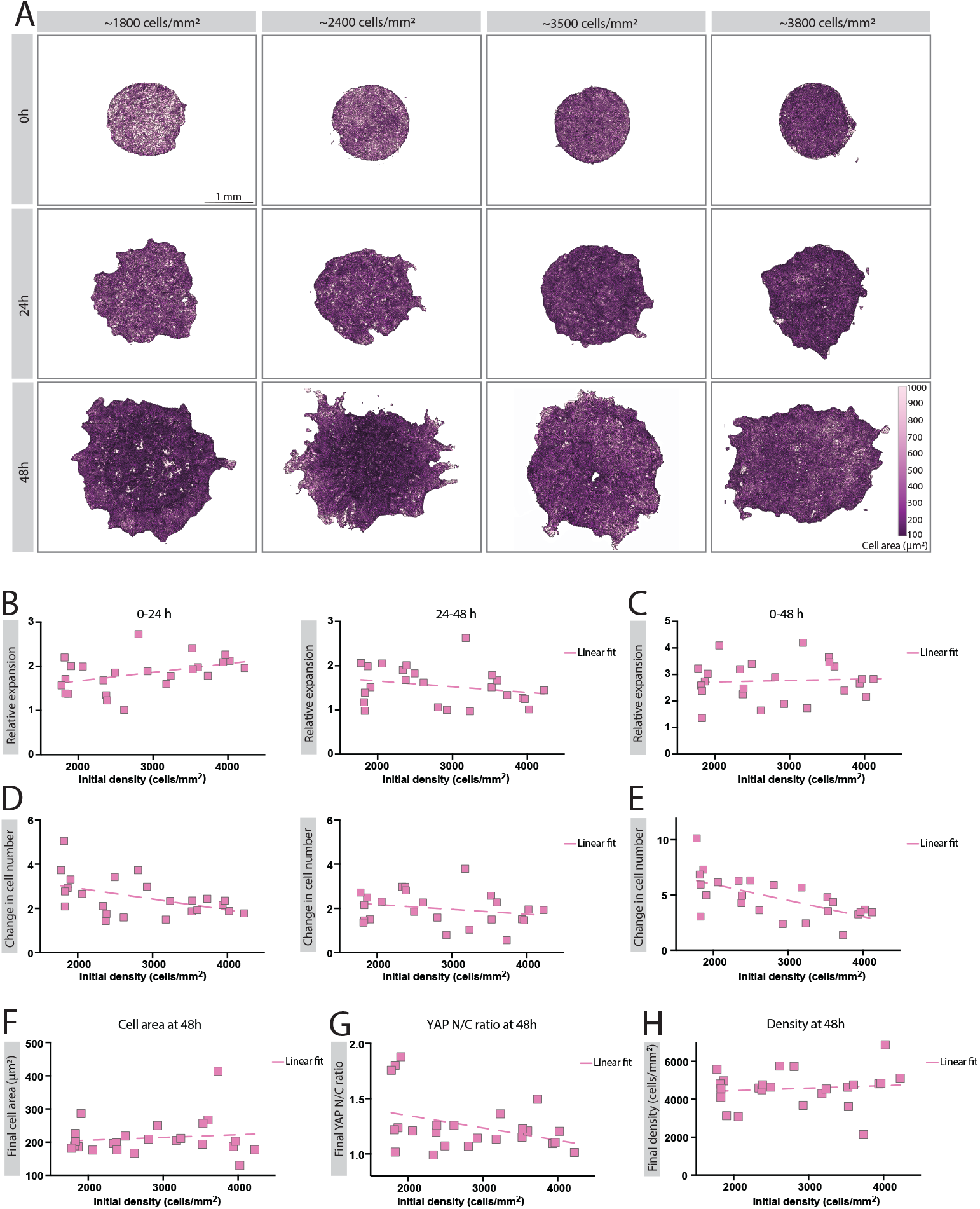
Tissue-wide cell number regulation couples epithelial expansion size to re-establishment of homeostasis. A: Cell-area colour maps of representative epithelial colonies with an initial diameter of 1.5 mm of varying densities ranging from ∼1800 to ∼3800 cells/mm^2^. Images acquired after 0, 24 and 48 hours of expansion, segmented and colour-coded according to cell area and with cell membrane outlines drawn as overlay. Scale bar, 1 mm. B: Relative expansion from 0-24, and 24-48 hours as a factor of initial density at 0h across the density range ∼1700 cells/mm^2^ to ∼4500 cells/mm^2^ (n = 24 colonies from at least 8 individual experiments). Relative expansion is defined as the fold change in colony area from start of expansion. Dashed line represents linear fit. C: Relative expansion from 0-48 hours as a factor of initial density at 0h across the density range ∼1700 cells/mm^2^ to ∼4500 cells/mm^2^ (n = 24 colonies from at least 8 individual experiments). Relative expansion is defined as the fold change in colony area from start of expansion. Dashed line represents linear fit. D: Change in cell number from 0-24, and 24-48 hours as a factor of initial density at 0h across the density range ∼1700 cells/mm^2^ to ∼4500 cells/mm^2^ (n = 24 colonies from at least 8 individual experiments). Dashed line represents linear fit. E: Change in cell number from 0-48 hours as a factor of initial density at 0h across the density range ∼1700 cells/mm^2^ to ∼4500 cells/mm^2^ (n = 24 colonies from at least 8 individual experiments). Dashed line represents linear fit. F: Final cell area at 48 hours as a factor of the initial density at 0 hours of colonies across the density range ∼1700 cells/mm^2^ to ∼4500 cells/mm^2^ (n = 24 colonies from at least 8 individual experiments). Dashed line represents linear fit. G: Final YAP N/C ratio at 48 hours as a factor of the initial density at 0 hours of colonies across the density range ∼1700 cells/mm^2^ to ∼4500 cells/mm^2^ (n = 24 colonies from at least 8 individual experiments). Dashed line represents linear fit. H: Final density at 48 hours as a factor of the initial density at 0 hours of colonies across the density range ∼1700 cells/mm^2^ to ∼4500 cells/mm^2^ (n = 24 colonies from at least 8 individual experiments). Dashed line represents linear fit.

In addition to quantifying tissue expansion at 24 and 48 hours, we measured changes in cell area, YAP N/C, and total cell number across the range of densities (Fig. 2B–G). We found that for the first 24 hours following confinement release, colonies originating from higher initial densities exhibited slightly greater tissue expansion, whereas colonies originating from lower initial densities showed a larger increase in total cell number over the same period, consistent with their higher YAP N/C at the time of release (Fig. 2B, Fig. 2D). During the subsequent 24 hours, differences in expansion across densities were modest, while colonies originating from lower initial densities continued to show a greater net increase in cell number which accompanied slightly higher expansion (Fig. 2B, Fig. 2D). As a result, despite differences in early expansion and cell number dynamics, colonies across the full initial density range reached comparable final extents of expansion after 48 hours, resulting in overall tissue expansion to be independent of initial density (Fig. 2C). In contrast, cumulative changes in cell number over the 48-hour period remained density dependent, with colonies starting at lower initial densities exhibiting larger net increases in cell number (Fig. 2E). Together, these observations indicate that epithelial colonies converge to a similar expanded size after confinement release, while differences in cell number dynamics persist as a function of initial density, reflecting density-dependent regulation of cell number rather than differences in overall tissue expansion. These results show that YAP activity in confined epithelial monolayers varies nonlinearly with cell density prior to expansion, defining distinct density ranges with different sensitivities to crowding. YAP activity is uniformly elevated at lower densities and decreases with increasing overall colony density, where at higher densities it is globally suppressed and only weakly sensitive to further crowding.

Given these density-dependent differences in cell number dynamics coupled with overall similar expansion size, we next examined whether epithelial colonies converged to a common cell-scale mechanochemical state after 48 hours of expansion. This analysis showed that despite significant starting density dependent differences in cell number, cell area, and YAP activity at the onset of expansion post confinement, all of these parameters converged to similar values after 48 hours, leading to an overall convergence of all expanding colonies to a final density of ∼4500 cells/mm^2^ (Fig. 2F-H).

Together, these results show that epithelial expansion is dynamically coupled to regulation of mechanochemical state and cell number, enabling robust, density-independent expansion and convergence to common homeostatic state.

### Agent-based modelling reveals mechanochemical feedback drives tissue-scale pressure regulation and density homeostasis in expanding epithelial colonies

To explore the mechanisms underlying cell number dynamics and convergence to a homeostatic state in expanding epithelial tissues post confinement, we next turned to computational modelling. Several prior models have coupled epithelial behaviour with proliferation dynamics and demonstrated that tissues can converge to these common states, using simple forms of mechanosensitivity such as coupling tissue growth rates to local pressure or density (15, 27, 29–33). While such frameworks, including models based on the Fisher-Kolmogorov equation and their mechanosensitive extensions, can exhibit convergence, they do not explicitly track the internal mechanochemical state of cells which we measured in our experiments. Thus, here we adopted a previously developed agent-based model that explicitly couples collective tissue mechanics to intracellular cell-cycle regulation and has been shown to reproduce key aspects of MDCK monolayer dynamics (34, 35). In this model, cells are represented as soft repulsive discs, and each cell contains a simplified cell-cycle oscillator, where proliferation and growth are governed by a cell cycle activity variable that is downregulated by isotropic mechanical compression, i.e. pressure, experienced by cells due to crowding (Fig. 3A, Model Suppl Note). This framework captures the non-instantaneous mechanochemical feedback between pressure and proliferation, conceptually analogous to experimentally observed density-dependent regulation of YAP activity and changes in cell numbers (8, 19) (Fig. 1B, 1C). This allows for history-dependent responses that cannot be captured by phenomenological, instantaneous coupling.

**Figure 3.**
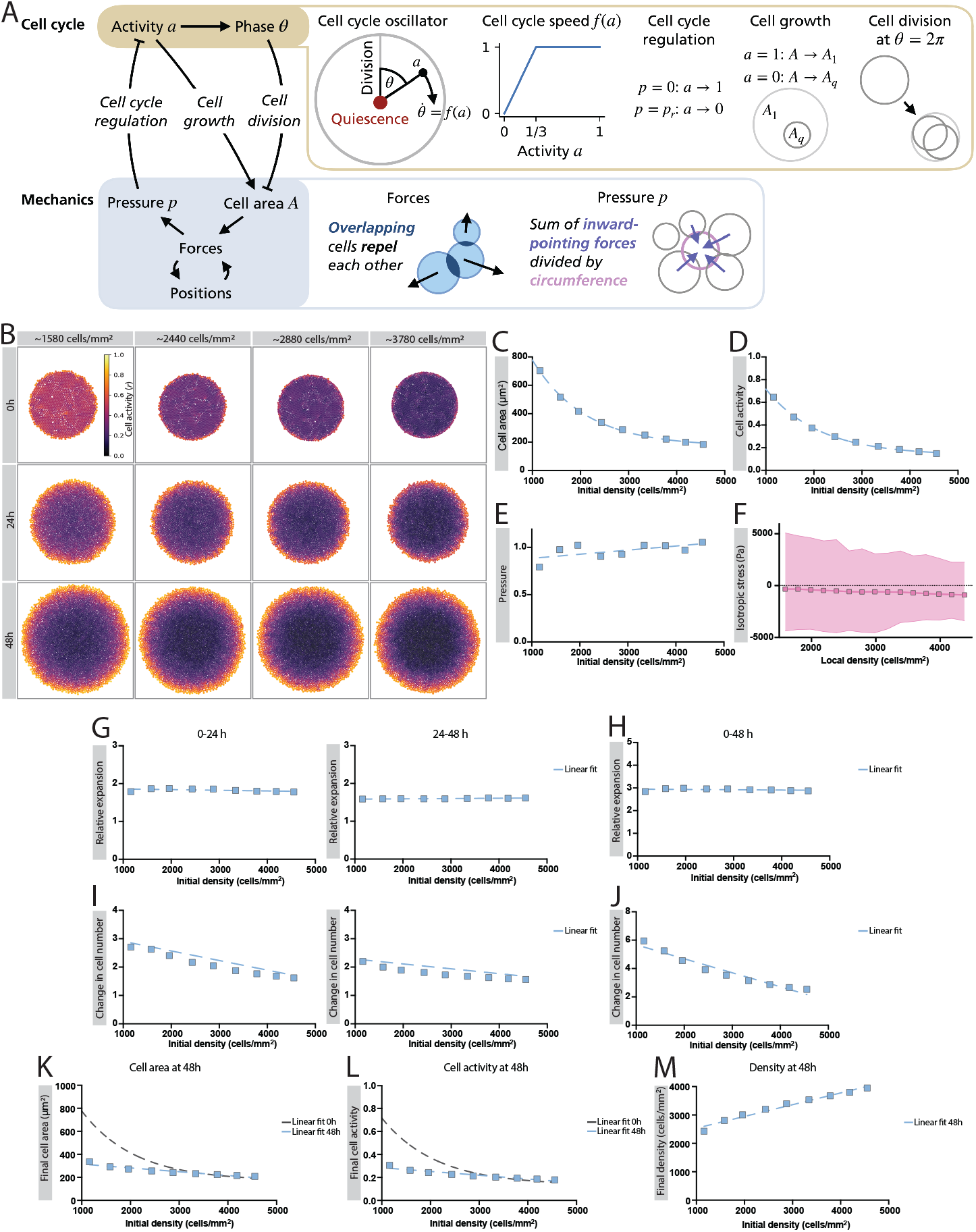
Agent-based modelling reveals mechanochemical feedback drives tissue-scale pressure regulation and density homeostasis in expanding epithelial colonies. A: Cartoon of the model: Each cell contains a stylized cell cycle oscillator with phase theta and activity r. Cells divide when the phase completes a full revolution, θ = n*2*π (with a natural number n). The activity 0 < 1 <= 1 regulates the speed of the cell cycle via a function f(r) (see small panel to the right): Cell divisions happen on average every 18h for r > ⅓ and decrease linearly with r for 0 < r < ⅓. B: Colour maps of simulations of epithelial colonies with an initial diameter of 1 mm of varying densities ranging from ∼1500 to ∼3800 cells/mm^2^ at 0, 24 and 48 hours of expansion. Colour-coded according to the model read-out cell activity value from 0-1. C: Cell area in µm^2^ as a factor of initial density at 0h across the density range ∼1500 cells/mm^2^ to ∼4500 cells/mm^2^ (n = 9 colonies from model simulations). D: Cell activity as a factor of initial density at 0h across the density range ∼1500 cells/mm^2^ to ∼4500 cells/mm^2^ (n = 9 colonies from model simulations). E: Pressure as a factor of initial density at 0h across the density range ∼1500 cells/mm^2^ to ∼4500 cells/mm^2^ (n = 9 colonies from model simulations). F: Isotropic stress (Pa) as a factor of the local density across density range of 1500 cells/mm^2^ to 4500 cells/mm^2^. Mean values represented with a solid connected line, whilst background points represent individual cells, shaded area represents full range of all measured cells at the given density. G: Relative expansion from 0-24, and 24-48 hours as a factor of initial density at 0h across the density range ∼1500 cells/mm^2^ to ∼4500 cells/mm^2^ (n = 9 colonies from model simulations). Relative expansion is defined as the fold change in colony area from start of expansion. Dashed line represents linear fit. H: Relative expansion from 0-48 hours as a factor of initial density at 0h across the density range ∼1500 cells/mm^2^ to ∼4500 cells/mm^2^ (n = 9 colonies from model simulations). Relative expansion is defined as the fold change in colony area from start of expansion. Dashed line represents linear fit. I: Change in cell number from 0-24, and 24-48 hours as a factor of initial density at 0h across the density range ∼1500 cells/mm^2^ to ∼4500 cells/mm^2^ (n = 9 colonies from model simulations). Dashed line represents linear fit. J: Change in cell number from 0-48 hours as a factor of initial density at 0h across the density range ∼1500 cells/mm^2^ to ∼4500 cells/mm^2^(n = 9 colonies from model simulations). Dashed line represents linear fit. K: Final cell area at 48 hours as a factor of the initial density at 0 hours of colonies across the density range ∼1500 cells/mm^2^ to ∼4500 cells/mm^2^ (n = 9 colonies from model simulations). Grey dashed line represents linear fit at 0h, and blue dashed line represents linear fit at 48h. L: Final YAP N/C ratio at 48 hours as a factor of the initial density at 0 hours of colonies across the density range ∼1500 cells/mm^2^ to ∼4500 cells/mm^2^ (n = 9 colonies from model simulations). Grey dashed line represents linear fit at 0h, and blue dashed line represents linear fit at 48h. M: Final density at 48 hours as a factor of the initial density at 0 hours of colonies across the density range ∼1500 cells/mm^2^ to ∼4500 cells/mm^2^ (n = 9 colonies from model simulations). Dashed line represents linear fit.

Since the key assumption of the model is that epithelial monolayers experience predominantly compressive stresses whose magnitude increases with cell density, we first decided to validate this experimentally. To do so, we quantified tissue-scale isotropic stresses in MDCK monolayers as a function of local cell density using monolayer stress microscopy (MSM) (36–39). This data showed that indeed, across an MDCK monolayer, isotropic stresses were predominantly negative indicating a compressive state, and that the magnitude of compressive stresses increased monotonically with increase in local cell density (Fig. 3F, Supplementary Fig. S3A, S3B).

Next, we simulated our expansion assays in the model by allowing colonies to grow from a single cell under circular confinement until they reached set target densities similar to the density range in the experiments (∼1200-4500 cells/mm^2^) (Model Suppl Note, Fig SN3B). Examination of the simulated colonies at stencil release prior to the onset of expansion revealed that similar to experimental results, median cell area of the colonies decreased with increase in overall colony density (Fig 3C). Notably, while the cell activity parameter did not exhibit discrete density-dependent regimes as observed for YAP N/C in the experiments, it decreased with increasing colony density following an exponential decay that gradually approached a plateau at higher densities (Fig 3D).

To understand these results further, we next examined how the mechanical states get established in the epithelial monolayer during confinement. Simulations revealed that as cells grew and packed the stencil, they gradually accumulated internal mechanical pressure (Model Suppl Note, Fig SN3A, SN3B). This pressure continued to increase until the colony reached a stable reference pressure, defined here as the equilibrium level of isotropic compression at which further proliferation becomes suppressed (Model Suppl. Note, Fig SN3B). At this reference pressure, cell activity dropped sharply, reflecting the onset of confinement-induced growth inhibition (Model Suppl. Note Fig. SN3A). Because colonies at higher densities remained under confinement longer than colonies at lower densities, they spent more time at this reference pressure, resulting in lower activity at the time of stencil release (Model Suppl. Note Fig. SN3A). Despite these differences in initial density and cell activity, however, all colonies exhibited similar internal pressure distributions at the moment of release, although the pressure was slightly increased with increasing density, which led to comparable upregulation of activity during early expansion (Fig. 3D, 3E).

Upon reaching the target density, the stencils were released, and the simulated colonies were allowed to expand for 48 hours. Results on overall expansion size showed that again similar to experiments, all colonies expanded to similar final size independent of initial density and the starting cell activity parameter. Together with our experimental result, these results indicate that overall epithelial expansion post confinement is independent of density and mechanochemical state at confinement (Fig. 3B, Fig 3F-I).

Following stencil release, simulations showed that initial differences in activity and cell size progressively diminished over time. Across the full range of initial densities, both cell area and cell activity evolved from being strongly dependent on initial density at the onset of expansion to exhibiting only a weak dependence by 48 hours (Fig. 3J, 3K). Colonies initiated at lower densities exhibited higher proliferation rates during the early phase of expansion, whereas colonies initiated at higher densities increased their proliferation more gradually, such that proliferation rates converged by 48 hours (Fig. 3H, 3I). Because of these coordinated dynamics, colonies expanded by comparable amounts and progressively approached a similar final density across the full range of starting conditions by 48 hours just as in our experimental observations (Fig. 3L).

Together, these results demonstrate that pressure-mediated feedback on proliferation and growth is sufficient to qualitatively reproduce the density-independent expansion and convergence of cell size and density observed during epithelial tissue spreading. This minimal model thus offers a mechanistic framework for how local mechanical compression can regulate global tissue behaviour, enabling robust coordination of homeostasis during dynamic epithelial remodelling.

### Reduction of internal pressure during confinement results in density-dependent changes in tissue expansion

Results from the model above suggest that pressure buildup during confinement and the cell’s ability to sense this pressure plays a central role in regulating proliferation and restoring homeostasis after expansion. Since our model tracks cell-cycle-dependent activity, it also allows us to examine how mechanical history influences subsequent growth dynamics. As such we next asked whether transiently altering how cells perceive pressure could shift their expansion behaviour or affect re-establishment of homeostasis. To this end, we introduced a reduced perceived pressure (RPP) condition in the model (Fig. 4A).

**Figure 4.**
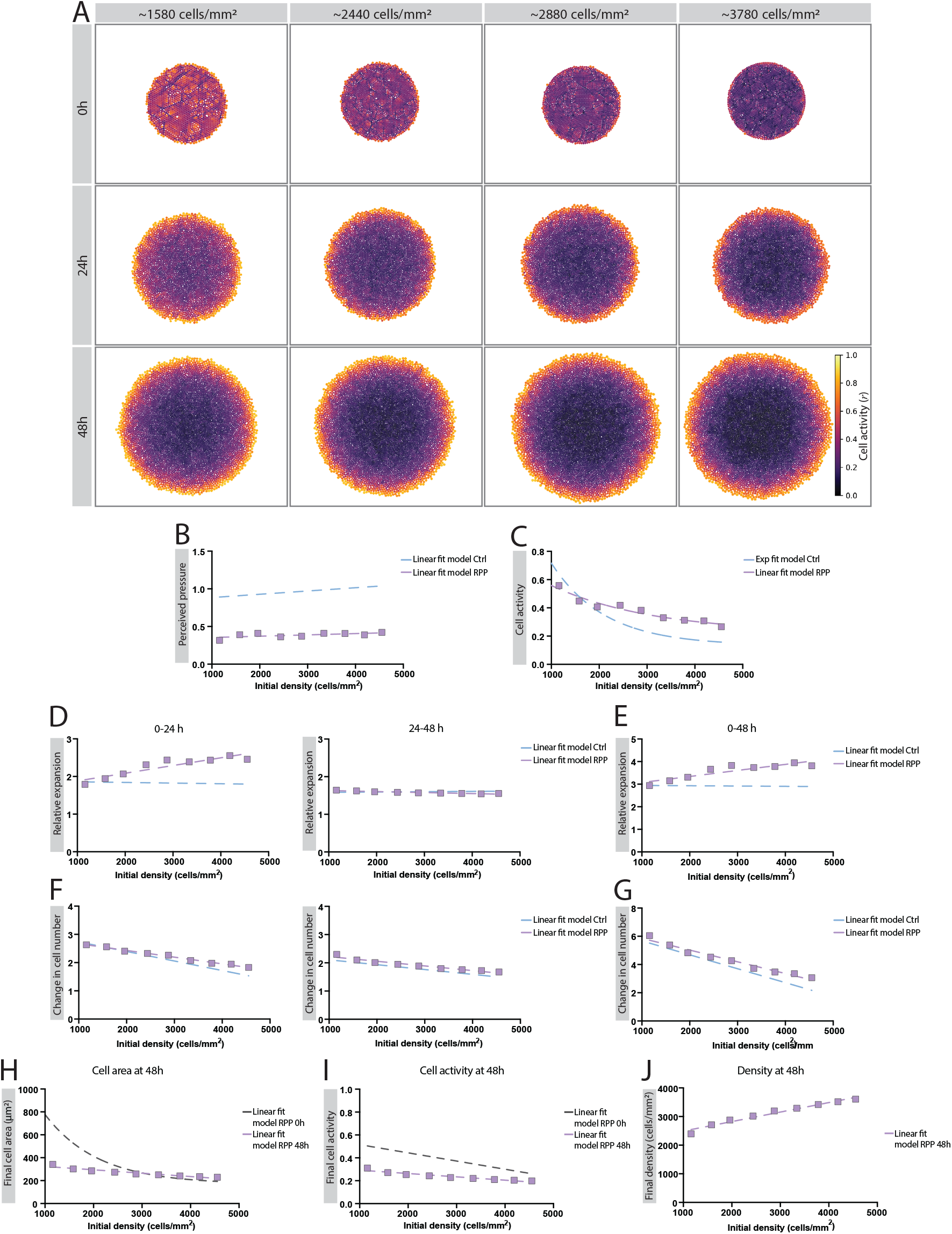
Reduction of internal pressure during confinement results in density-dependent changes in tissue expansion. A: Colour maps of RPP condition simulations of epithelial colonies with an initial diameter of 1 mm of varying initial densities ranging from ∼1500 to ∼3800 cells/mm^2^ at initiation of simulation (0h) and after 24 and 48 hours of expansion. Colour-coded according to the model read-out cell activity value from 0-1. B: Perceived pressure as a factor of initial density at initiation of experiments (0h) across the density range ∼1500 cells/mm^2^ to ∼4500 cells/mm^2^ for control and reduced perceived pressure (RPP) colonies (n = 9 colonies from model simulations). Purple dashed line representing the linear fit of RPP colonies and blue dashed line represents the linear fit of Ctrl colonies. C: Cell activity as a factor of initial density at initiation of experiments (0h) across the density range ∼1500 cells/mm^2^ to ∼4500 cells/mm^2^ for control and reduced perceived pressure (RPP) colonies (n = 9 colonies from model simulations). Purple dashed line representing the non-linear fit of RPP colonies and blue dashed line represents the non-linear fit of Ctrl colonies. D: Relative expansion from 0-24, and 24-48 hours as a factor of initial density at 0h across the density range ∼1500 cells/mm^2^ to ∼4500 cells/mm^2^ (n = 9 colonies from model simulations). Relative expansion is defined as the fold change in colony area from start of expansion. Dashed line represents linear fit. E: Relative expansion from 0-48 hours as a factor of initial density at 0h across the density range ∼1500 cells/mm^2^ to ∼4500 cells/mm^2^ (n = 9 colonies from model simulations). Relative expansion is defined as the fold change in colony area from start of expansion. Purple dashed line representing the linear fit of RPP colonies and blue dashed line represents the linear fit of Ctrl colonies. F: Change in cell number from 0-24, and 24-48 hours as a factor of initial density at 0h across the density range ∼1500 cells/mm^2^ to ∼4500 cells/mm^2^ (n = 9 colonies from model simulations). Purple dashed line representing the linear fit of RPP colonies and blue dashed line represents the linear fit of Ctrl colonies. G: Change in cell number from 0-48 hours as a factor of initial density at 0h across the density range ∼1500 cells/mm^2^ to ∼4500 cells/mm^2^ (n = 9 colonies from model simulations). Purple dashed line representing the linear fit of RPP colonies and blue dashed line represents the linear fit of Ctrl colonies. H: Final cell area at 48 hours as a factor of the initial density at 0 hours of RPP colonies across the density range ∼1500 cells/mm^2^ to ∼4500 cells/mm^2^ (n = 9 colonies from model simulations). Grey dashed line represents linear fit at 0h, and purple dashed line represents linear fit at 48h. I: Final YAP N/C ratio at 48 hours as a factor of the initial density at 0 hours of RPP colonies across the density range ∼1500 cells/mm^2^ to ∼4500 cells/mm^2^ (n = 9 colonies from model simulations). Grey dashed line represents linear fit at 0h, and purple dashed line represents linear fit at 48h. J: Final density at 48 hours as a factor of the initial density at 0 hours of RPP colonies across the density range ∼1500 cells/mm^2^ to ∼4500 cells/mm^2^ (n = 9 colonies from model simulations). Dashed line represents linear fit.

Here, we selectively dampened the mechanical signal sensed by cells, without altering the actual tissue-level pressure (Supplementary Fig. S4A). Specifically, we lowered the perceived pressure by 60% for a 3-hour period immediately before stencil release across the full range of initial densities, while disabling cell growth and division during this time to isolate the effect (Fig. 4B). Surprisingly, this brief reduction in perceived pressure had a strong, density-dependent effect on cell activity (Fig. 4C). Colonies initiated at lower densities, where activity was already near maximal, showed little response to RPP, whereas colonies initiated at higher densities exhibited a marked increase in activity relative to control (Fig 4C). This reflects the fact that colonies experiencing stronger confinement-induced suppression prior to release respond more strongly to transient relief of perceived pressure, restoring activity that would otherwise remain low. At the time of confinement release, RPP-treated colonies exhibited elevated activity compared to control colonies of comparable density, demonstrating that cellular responses to pressure depend on prior mechanical history rather than initial density alone.

Following stencil release, we reinstated cell growth and division and allowed colonies to expand freely for 48 hours (Fig. 4A). During this phase, the RPP condition was gradually reversed and perceived pressure sensing recovered exponentially toward the model-defined reference pressure (Supplementary Fig. S4B). Despite this restoration, the early colony density-dependent change in activity induced by RPP had lasting consequences; during the first 24 hours, the rate of expansion increased with initial density, driven by a sustained increase in proliferation (Fig. 4D-G). Strikingly, this enhanced expansion in RPP colonies with higher initial density acted as a compensatory response to the early surge in proliferation, allowing the tissue to regulate its density and converge with lower density colonies showing less expansion over time. As a result, despite starting fr increasing initial density, similar to control colonies from distinct mechanochemical states, all colonies ultimately started converging to similar final cell densities by 48 hours with cell activity and average cell sizes also equilibrated across the density range conditions after 48 hours of expansion (Fig. 4J-L).

Together, these results show that transient modulation of pressure sensing during confinement alters early expansion dynamics in a density-dependent manner. Colonies subjected to reduced perceived pressure exhibited enhanced early expansion at higher densities, while all conditions ultimately converged to a common final density. Thus, transient mechanical perturbations reshape expansion trajectories without disrupting robust restoration of tissue homeostasis.

### Transient mechanical resetting during confinement selectively elevates YAP activity at higher densities

To experimentally test the model’s prediction that transient reductions in perceived mechanical pressure can modulate cell numbers in a density-dependent manner, we sought an approach to mimic the RPP condition used in simulations. In the model, RPP temporarily decouples the isotropic stresses experienced by cells under confinement from the mechanotransduction signal that regulates their activity, revealing a latent capacity for high-density tissues to re-activate and expand. Since cells in epithelial tissues sense pressure indirectly through local mechanical stresses, such as cortical tension and junctional forces, we used a blebbistatin (Bb) washout approach. Bb, a myosin II inhibitor that reduces actomyosin contractility and lowers cortical tension in epithelial cells, has been shown to decrease cell-generated stresses and impair mechanosensing in MDCK epithelial monolayers (16, 37, 40–44). Although Bb affects multiple aspects of cell physiology, its most direct and well-established effect in this context is the transient reduction of actomyosin-generated mechanical stresses (37, 40, 44, 45). To verify the effectiveness and transient nature of Bb washout, we assessed cytoskeletal organization and tissue-scale mechanics in confluent MDCK epithelial monolayers following Bb treatment and washout. As expected, actin fibres were no longer visible across the epithelial monolayer consistent with loss of cortical actin and isotropic stresses after 1 hour of 50 µM Bb treatment (Supplementary Fig. S5A). We then washed out the Bb and added fresh media and could observe recovery of stress fibres within the first 15 minutes (Supplementary Fig. S5A). By 30 minutes of washout, actin fibres had completely recovered to levels comparable to untreated controls (Supplementary Fig. S5A). To then verify functional recovery, we performed MSM and found that 30 minutes after washout was sufficient for recovery of mean isotropic stress to near-control levels, while maintaining the normal density-dependent scaling of compressive stress across the monolayer (Supplementary Fig. S5B, S5C). Notably, Bb washout also resulted in a significant reduction in stress heterogeneity across the monolayer, indicating a more mechanically homogeneous tissue state (Supplementary Fig. S5C). Together with previous studies showing that Bb treatment reduces isotropic stresses in MDCK monolayers (37, 45), these results demonstrate that transient Bb treatment followed by washout induces a uniform mechanical reset at the tissue scale, without altering epithelial integrity or cell density.

After confirming the effect of Bb washout (Bb WO), epithelial monolayers were grown in PDMS stencils across the density range just as before and treated with 50 µM Bb for 1 hour upon reaching target densities, washed out and then allowed to recover for 30 minutes prior to stencil release. Colonies were then fixed at the time of stencil removal and analyzed for cell area and YAP N/C to quantify the mechanochemical state prior to onset of expansion (Fig. 1A). Analysis of cell area across all colonies showed that median cell area in Bb WO colonies still decreased monotonically with increasing initial density, similar to control colonies in Fig. 1 (Fig 5A, 5B). YAP N/C in Bb WO colonies also showed density dependence in Bb WO colonies though with differences compared to control colonies (Fig 5C). Bb WO led to selective increase in YAP N/C at higher densities while having minimal effect at lower densities,resulting in a flatter YAP N/C– density relationship compared to control (Fig. 5A, 5C). This density-dependent upward shift in YAP N/C closely mirrored the change in cell activity predicted by the model under RPP conditions. To directly compare the effect across densities with controls in Fig 1, colonies were grouped by initial density just as before. In colonies <2500 cells/mm^2^, YAP N/C values in Bb WO conditions were not significantly different from those observed in control colonies. In contrast, Bb WO colonies >2500 cells/mm^2^ exhibited a significant increase in YAP N/C (Fig. 5A, 5D). Spatial analysis further showed that this increase in Bb WO colonies >2500 cells/mm^2^ reflected a population-wide shift toward more nuclear YAP, with high YAP N/C cells homogeneously distributed throughout the colony rather than restricted to specific regions or colony edge (Fig. 5E).

**Figure 5.**
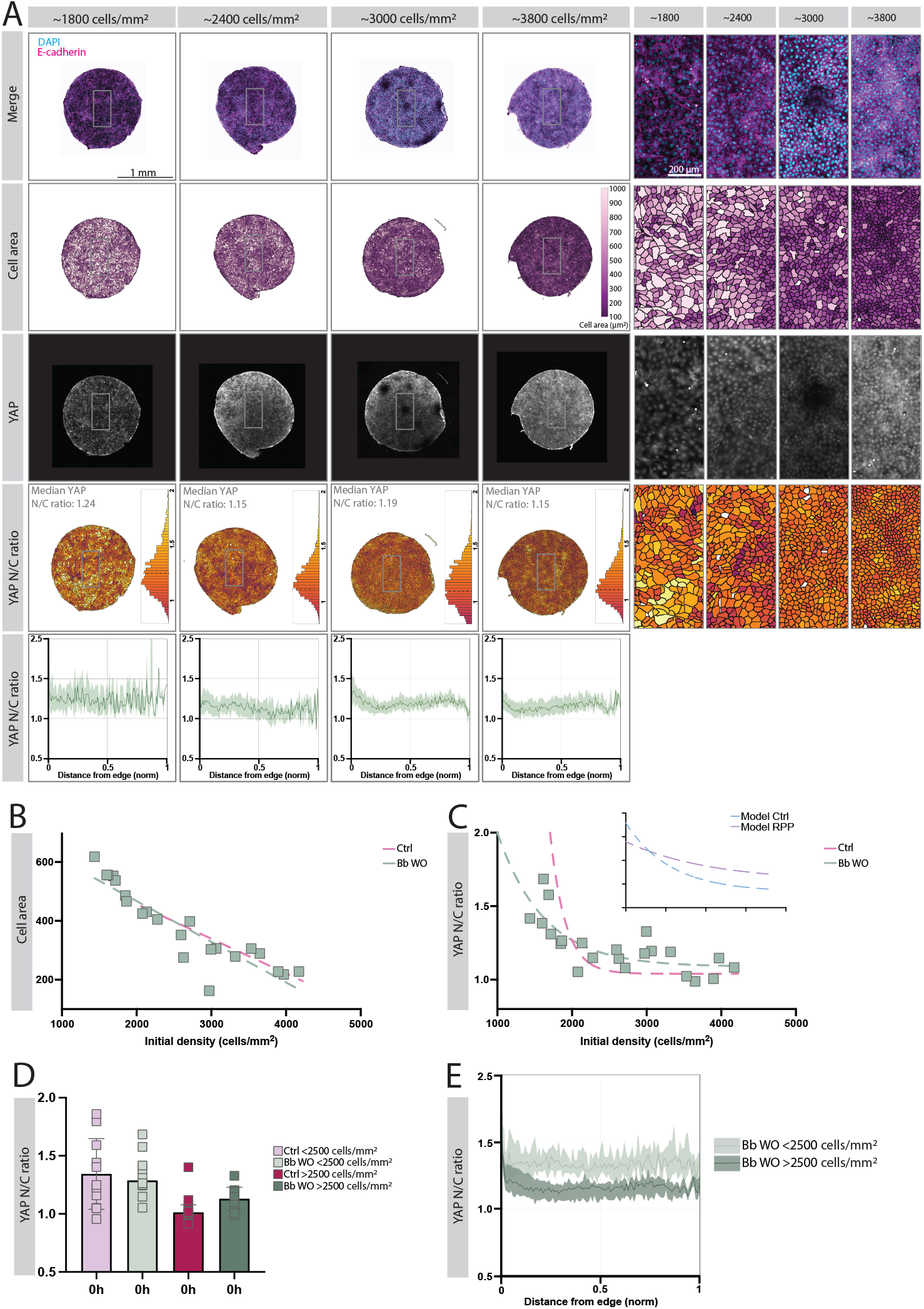
Transient mechanical resetting during confinement selectively elevates YAP activity at higher densities. A: Representative images of DAPI/E-cadherin merge, cell area colour maps, YAP channel and YAP N/C colour maps of colonies of varying densities ranging from ∼1800 to ∼3800 cells/mm^2^ after Bb WO treatment. Images taken after removal of stencil, color-coded for cell area and YAP N/C ratio. Representative histograms display the population spread for YAP N/C ratios and colour scale from 0.8-2. The dashed lines indicate the median YAP N/C ratio for the colony. Radial spatial plots displaying median YAP N/C ratio of example colonies. Thicker line represents median YAP across colony, with shaded error bars representing interquartile range (IQR) of these. X-axis represents the distance from edge to centre, that has been normalized to arbitrary values. Scale bar whole colony images, 1 mm. Scale bar zoom ins 200 µm. B: Cell area in µm^2^ as a factor of initial density at 0h across the density range ∼1400 cells/mm^2^ to ∼4200 cells/mm^2^ after Bb WO treatment (n = 22 colonies from at least 6 individual experiments). Green dashed line represents linear fit of Bb WO colonies and pink dashed line represents linear fit of Ctrl colonies. C: YAP N/C ratio as a factor of initial density at 0h across the density range ∼1700 cells/mm^2^ to ∼4500 cells/mm^2^ after Bb WO treatment. (n = 22 colonies from at least 6 individual experiments). Dashed line in green represents non-linear fit of Bb WO colonies, dashed line in pink represents non-linear fit of Ctrl colonies. Inset showing non-linear fits of model ctrl (blue) and model RPP (purple). D: YAP N/C ratio of control and Bb WO colonies <2500 and >2500 cells/mm^2^ imaged and measured after removal of stencil (n = 11 <2500 cells/mm^2^ and 13 >2500 cells/mm^2^ colonies from at least 8 individual experiments, and 10 LD Bb <2500 cells/mm^2^ and 12 Bb WO >2500 cells/mm^2^ colonies from at least 6 individual experiments). E: Radial spatial plots displaying median YAP N/C ratio of Bb WO colonies, divided into colonies with a density <2500 cells/mm^2^ and >2500 cells/mm^2^. Thicker line represents median YAP ratio across colony, with shaded error bars representing interquartile range (IQR) of these. X-axis represents the distance from edge to centre, that has been normalized to arbitrary values

Together, these results show that Bb WO acts as a transient mechanical reset of epithelial colony during confinement. This reset selectively alters the mechanochemical state of higher-density colonies, elevating YAP activity in a manner that closely mirrors the RPP condition in the model.

### Resetting stresses during confinement leads to changes in density-dependent tissue expansion

We next allowed Bb WO colonies to expand for 24- and 48-hours following stencil release before fixation and analysis of expansion and changes in cell number (Fig. 6A). Compared to control colonies, Bb WO colonies exhibited marked differences in expansion at both time points (Fig. 6B–C). During the first 24 hours after release, Bb WO colonies expanded more than controls across the full range of initial densities, with this effect being most pronounced in colonies initiated at higher densities (Fig. 6B). During the subsequent 24 hours, this enhancement in expansion was no longer apparent at lower initial densities, whereas colonies initiated at higher densities maintained a modestly elevated expansion rate relative to controls (Fig. 6B, 6C). As a result, Bb WO colonies displayed a clear initial density dependence in cumulative expansion over 48 hours, with higher-density colonies expanding significantly more than lower-density Bb WO colonies and control colonies (Fig. 6C). This enhanced expansion in higher density Bb WO colonies was accompanied by a selective, density-dependent increase in cell number (Fig. 6D, 6E). Consequently, net changes in cell number in Bb WO colonies became largely independent of initial density over the 48-hour expansion period, in contrast to the density dependence observed in control colonies (Fig. 6E).

**Figure 6.**
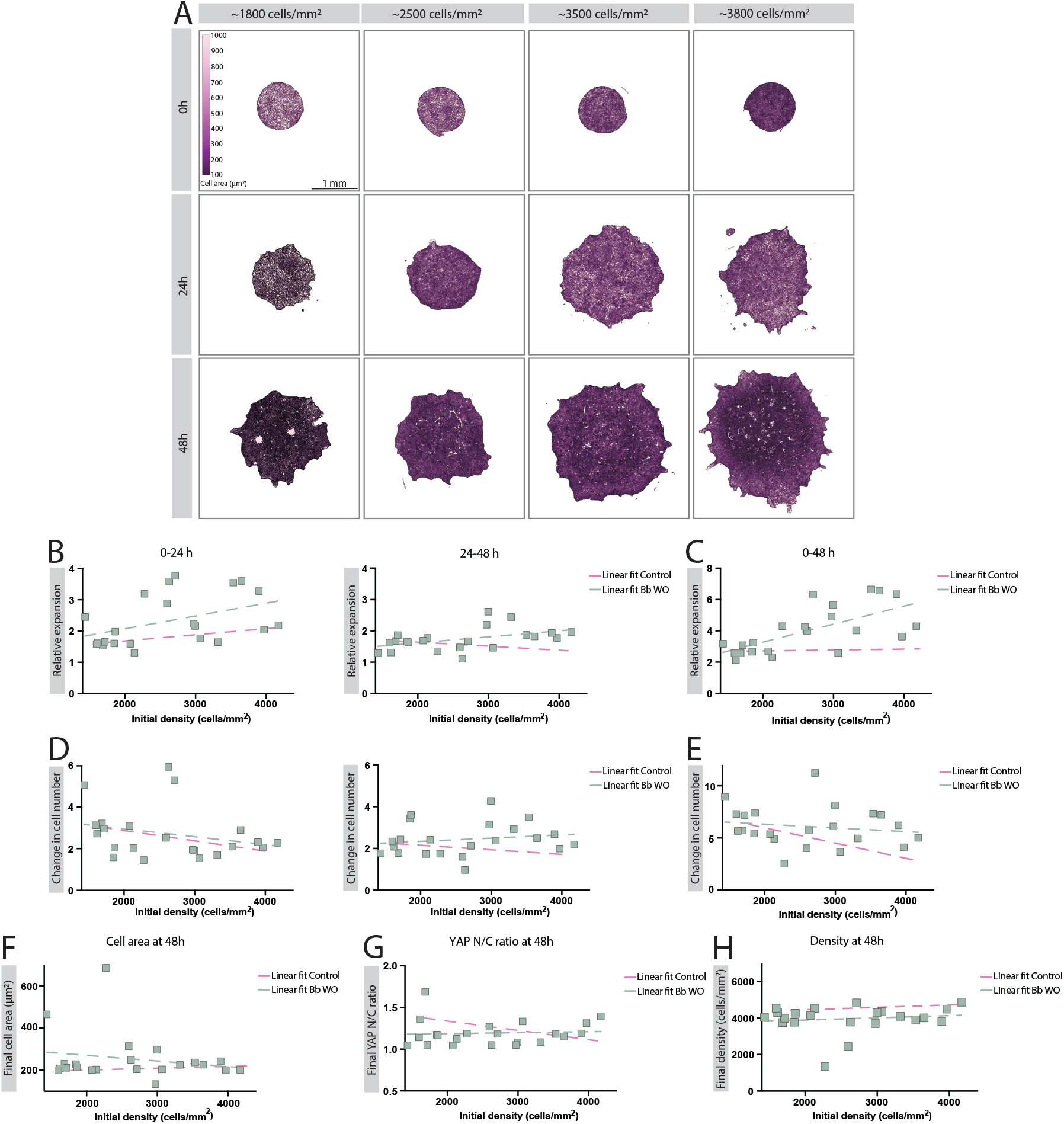
Resetting stresses during confinement leads to changes in density-dependent tissue expansion. A: Cell-area colour maps of representative epithelial colonies with an initial diameter of 1.5 mm of varying densities ranging from ∼1800 to ∼3800 cells/mm^2^ after Bb WO treatment. Images acquired after 0, 24 and 48 hours of expansion, segmented and colour-coded according to cell area and with cell membrane outlines drawn as overlay. Scale bar, 1 mm. B: Relative expansion from 0-24, and 24-48 hours as a factor of initial density at 0h across the density range ∼1400 cells/mm^2^ to ∼4500 cells/mm^2^ after Bb WO treatment (n = 22 colonies from at least 6 individual experiments). Dashed line in green represents linear fit of Bb WO colonies, dashed line in pink represents linear fit of control colonies. C: Relative expansion from 0-48 hours as a factor of initial density at 0h across the density range ∼1400 cells/mm^2^ to ∼4500 cells/mm^2^ Bb WO treatment (n = 22 colonies from at least 6 individual experiments). Dashed line in green represents linear fit of Bb WO colonies, dashed line in pink represents linear fit of control colonies. D: Change in cell number from 0-24, and 24-48 hours as a factor of initial density at 0h across the density range ∼1400 cells/mm^2^ to ∼4500 cells/mm^2^ Bb WO treatment (n = 22 colonies from at least 6 individual experiments). Dashed line in green represents linear fit of Bb WO colonies, dashed line in pink represents linear fit of control colonies. E: Change in cell number from 0-48 hours as a factor of initial density at 0h across the density range ∼1400 cells/mm^2^ to ∼4500 cells/mm^2^ Bb WO treatment (n = 22 colonies from at least 6 individual experiments). Dashed line in green represents linear fit of Bb WO colonies, dashed line in pink represents linear fit of control colonies. F: Final cell area at 48 hours as a factor of the initial density at 0 hours of colonies across the density range ∼1400 cells/mm^2^ to ∼4500 cells/mm^2^ after Bb WO treatment (n = 22 colonies from at least 6 individual experiments). Dashed line in green represents linear fit of Bb WO colonies, dashed line in pink represents linear fit of control colonies. G: Final YAP N/C ratio at 48 hours as a factor of the initial density at 0 hours of colonies across the density range ∼1400 cells/mm^2^ to ∼4500 cells/mm^2^ after Bb WO treatment (n = 22 colonies from at least 6 individual experiments). Dashed line in green represents linear fit of Bb WO colonies, dashed line in pink represents linear fit of control colonies. H: Final density at 48 hours as a factor of the initial density at 0 hours of colonies across the density range ∼1400 cells/mm^2^ to ∼4500 cells/mm^2^ after Bb WO treatment (n = 22 colonies from at least 6 individual experiments). Dashed line in green represents linear fit of Bb WO colonies, dashed line in pink represents linear fit of control colonies.

Despite these altered expansion trajectories, analysis across the full range of initial densities revealed that Bb WO colonies converged to the final density and YAP N/C ratio by 48 hours, indicating preservation of density homeostasis (Fig. 6F, 6G). Average cell area also converged across conditions, confirming that differences in final colony size reflected differences in cell number rather than sustained changes in cell size (Fig. 6H). To further facilitate direct comparison with control experiments and all model predictions, colonies were again grouped based on the YAP N/C–defined density regimes in control colonies. In this binned analysis, Bb WO colonies with initial densities >2500 cells/mm^2^ exhibited significantly greater increase in colony area and cell number compared to all other conditions, while colonies <2500 cells/mm^2^ showed no difference (Supplementary Fig. S6A–D). Across all conditions, binned analyses confirmed convergence of final density, cell area, and YAP activity, consistent with the experimentally observed and model-predicted approach to a common homeostatic state (Supplementary Fig. S6E-J).

Together, these results demonstrate that transient reduction of crowding-induced pressure sensing during confinement selectively alters density-dependent population growth in high-density tissues. This confirms the model’s prediction that epithelial expansion is regulated by time-integrated mechanical stress and mechanical memory, rather than by instantaneous density alone.

## Discussion

In this study, we provide an experimental demonstration and theoretical rationalization of a reversible, history-dependent pressure-sensing mechanism that links molecular activity, cell-cycle dynamics, and collective tissue expansion. In agreement with earlier studies, epithelial monolayers starting from a wide range of initial densities expand to similar final sizes and densities, revealing an emergent capacity for tissue-scale self-regulation. Here we show that this convergence occurs despite pronounced differences in initial YAP activity, cell size, and cell number dynamics, indicating that expansion is governed by dynamically regulated collective behaviour rather than initial conditions alone. A mechanochemical model that incorporates pressure-mediated feedback on cell-cycle activity recapitulates these dynamics. In the model, pressure builds during confinement, modulates progression through the cell cycle, and thereby regulates cell number accumulation, leading to convergence in density. Transient perturbation of this pressure-mediated feedback, either in silico by reducing perceived pressure or in vitro via blebbistatin washout, selectively restores cell-cycle activity selectively in higher-density tissues while preserving the final homeostatic state. Together, these results support a framework in which the internal state of cells integrates local mechanical stress and crowding over time, coordinating collective growth to robustly maintain epithelial homeostasis.

Previous work has shown that epithelial tissues can maintain functional integrity despite local heterogeneity in signalling and morphology, and that confinement can modulate growth through mechanical feedback (7, 16, 25, 29, 30, 46–49). Such behaviour is well captured by continuum and mean-field frameworks, including Fisher–Kolmogorov–type equations and their mechanosensitive extensions, in which growth rates decrease with increasing density or pressure. These models also successfully explain how tissues can average out local variability and converge toward a shared target state. However, most of these approaches assume an instantaneous coupling between mechanical signals and cellular responses. Our findings indicate that this assumption is insufficient to capture the full range of tissue-scale behaviours observed during expansion. Instead, epithelial cells retain an internal dynamic state, here reflected in cell-cycle activity, that integrates mechanical inputs over time and causally modulates collective expansion behaviour. As a result, transient perturbations of pressure sensing can produce pronounced, density-selective differences in early expansion dynamics even when boundary conditions and initial densities are identical. Importantly, despite these divergent early trajectories, tissue-scale homeostasis remains robust, with coordinated expansion restoring pressure balance and enabling convergence to a common final density.

Central to this process appears to be a set of mechanosensitive regulators of cell activity, including, but not limited to, the transcriptional co-activator YAP. In our system, YAP activity was spatially uniform and dynamically modulated by both initial density and changes in mechanical stress. Similar roles have been reported for other mechanosensitive regulators, including β-catenin, Piezo1, and cyclin D1, in controlling stretch- or density-dependent cell-cycle progression (19, 20, 22, 47). These factors may act in parallel or downstream of shared mechanical signals, forming a distributed control system that links tissue-scale mechanics to cell-cycle regulation. Such a separation between mechanical sensing and downstream execution may enable flexible yet robust control of growth and division under dynamic conditions. Consistent with this view, the selective restoration of activity in higher-density colonies following transient stress reduction supports the existence of an integrated, feedback-driven mechanism governing collective tissue growth.

Our modelling and blebbistatin experiments suggest that total pressure, emerging from confinement and cumulative mechanical stress, acts as a mechanical signal that is sensed by cells and incorporated into their internal regulatory state. Colonies that experienced prolonged high pressure exhibited suppressed cell-cycle activity and reduced cell number accumulation, whereas transient mechanical relaxation reversed this suppression and restored growth capacity, without altering the final density achieved. This points to a feedback mechanism in which cells integrate mechanical history through their internal state to regulate growth and division dynamics, resembling checkpoint-like behaviour. Similar mechanically regulated checkpoints have been described in other systems, including tension-sensitive G2 arrest via E-cadherin–Wee1 signalling, as well as β-catenin– or cyclin D1–mediated responses to crowding, suggesting that such feedback loops may be a general feature of mechanically regulated tissues (16, 19, 47, 50). Together, these findings extend existing frameworks of mechanical control by explicitly incorporating cellular memory and internal state.

We note that recent models such as Höllring et al. (2023) and Höllring et al., PNAS (2024) represent important advances in mechanosensitive tissue modelling (15, 30). Whether such frameworks can reproduce the full set of history-dependent responses revealed by transient mechanical resetting remains to be tested. In this context, our study provides an experimental benchmark and theoretical rationale for a reversible, history-dependent pressure-sensing mechanism that links molecular activity, cell-cycle dynamics, and collective tissue expansion.

## Methods

### Cell culture

E-cadherin-RFP expressing Madin-Darby Canine gII cells (MDCKs) were cultured in DMEM Low glucose (Gibco, 31885023) supplemented with 10% FBS (Gibco, 10270106), 1 g/l sodium bicarbonate (Sigma, 144-55-8), 500 ug/ml G-418 solution (Roche, 4727878001) and penicillin/Streptomycin (Gibco, 15140122) at 37°C and 5% CO_2_. Cells were used for a maximum of 20 passages.

### Cell seeding

50 mm^2^ glass coverslips were coated with 10 µg/ml of Fibronectin (FN) (Sigma Aldrich, 0895) at 37°C for 30 minutes. Coverslips were then rinsed in PBS and PDMS stencils were placed on top and gently pressed down to adhere. MDCK-E-cadherin-RFP cells were seeded at appropriate cell numbers for desired density within the PDMS wells and left to attach for 30-60 minutes before wells were flooded with DMEM. Cells were left to expand at 37°C and 5% CO_2_ overnight. The PDMS stencils were peeled off and cells were fixed according to the method described below either 10 minutes after peeling (for the 0 hour timepoint) or placed back in the incubator with fresh media for an additional 24 or 48 hours prior to fixation (for the 24- and 48-hour timepoints).

### Immunostaining

Cells were fixed with 4% formaldehyde (Thermo scientific, 28906) in phospho-buffered saline (PBS) for 15 min at room temperature (RT). After fixation, cells were permeabilised with 0.2% Triton X (Alfa Aesar, A16046) in PBS for 5 min at RT. Cells were rinsed with PBS, and then blocked with 3% BSA (Sigma-Aldrich, A7906) in PBS at RT for at least 45 minutes, or at 4°C overnight. For analysis of YAP, cells were incubated with primary mouse anti-YAP monoclonal antibody (1:100) (Santa Cruz, sc-101199) in 1% BSA in PBS overnight at 4°C. Cells were washed with 0.05% Tween (Fischer Scientific, BP337-100) in PBS 3 times 1 minutes with gentle rocking and then incubated with 1:400 secondary Alexa Fluor 488 goat anti-mouse Ab, for cell count using nuclear signal 1:500 Hoechst 33342 (Thermo Scientific, 62249) in 1% BSA in PBS was added with the secondary Ab for 1 hour at RT prior to mounting the coverslips on slides for imaging.

### Confocal microscopy

Images were acquired using a 20x (0.75 NA)(Plan APO) objective on a Nikon Confocal A1RHD microscope, with 405-, 488-, and 561-nm laser lines and equipped with GaAsP PMT and PMT detectors. To capture the full cell volume z-stacks with a step size of 1.5mm were acquired. A custom Nikon Jobs script was used to acquire multi-point z-stacks across the whole colony.

### Blebbistatin wash-out

Cells were seeded on coverslips as previously described, and upon reaching desired density media was replaced with media containing 50μM of Blebbistatin (Merck, 203390) in PBS. After 60 minutes of Bb incubation, cells were rinsed in PBS and kept for a further 20 minutes in DMEM at 37°C prior to peeling the stencils and performing the experiments as previously described.

### Image analysis

The image analysis was carried out using Julia (version 1.10.4) (51). Each fluorescence channel, E-cadherin, nuclear, and YAP, was reduced to a 2D image via maximum intensity projection. The membrane channel was used as a reference for stitching partially overlapping fields-of-view. For segmentation, a global threshold (Rosin’s method) was first applied to identify the stencil foreground, and adaptive thresholding with multiple window sizes was then used on the nuclear channel to separate nuclei (52). After thresholding, morphological filters and a watershed step were applied to improve the segmentation and separate adjacent nuclei. To segment the cytoplasmic region, the E-cadherin channel was used for thresholding, followed by a watershed algorithm to delineate each cell’s boundary around its nucleus (53). Duplicate cells detected in overlapping regions were merged so that each cell was only counted once. To quantify YAP localization, nuclear YAP intensity was measured directly, while cytoplasmic YAP was calculated in a perinuclear annulus defined by a distance transform, selecting pixels within a set range from the nucleus. The mean intensity in each mask was used to compute a nuclear-to-cytoplasmic intensity ratio for each cell.

### Monolayer stress microscopy

Polydimethylsiloxane (PDMS) substrates with elastic moduli of 3 kPa and 15 kPa were prepared by mixing the components A and B of CY 52-276 (Dow Corning) at ratios of 6:5 and 1:1, respectively (54). The mixture was spread into 35-mm Petri dishes and cured at 80 °C for 2 hours. The cured gels were silanized with 10% (3-aminopropyl)triethoxysilane (APTES, Sigma-Aldrich) in absolute ethanol for 10–15 minutes, rinsed thoroughly with absolute ethanol three times, and air-dried. For traction force microscopy, substrates were coated with carboxylate-modified fluorescent microspheres (0.2 µm, Invitrogen™ F8810) diluted 1:500 in Mq water. The bead solution was sonicated for 5 minutes before application. Following a 5-minute incubation, substrates were rinsed with Mq water three times, dried, and coated with 10 µg/ml human plasma fibronectin (in PBS) for 1–2 hours at 37 °C. Cells were seeded onto the functionalized substrates and cultured overnight to form a confluent monolayer prior to imaging.

Monolayers were imaged live on an inverted wide-field fluorescence microscope (Nikon Ti2-E) equipped with a 10× Plan Apo objective (NA 0.45), a CMOS camera (Nikon DS-Qi2), and an environmental chamber (Okolab) maintaining 37 °C and 5% CO_2_. Phase-contrast images of cells and wide-field fluorescence images of embedded microspheres (filter cube: Ex. 543-566 nm, Em. 582-636 nm) were acquired simultaneously every 10 minutes. At the end of each timelapse, cells were detached by adding 200 µL of 10% Triton X-100 to the medium, and a reference fluorescence image of the relaxed bead layer was acquired. Fluorescent bead images were processed using FIJI for stabilization and illumination correction. Bead displacement fields were calculated by registering each timelapse frame against the reference frame using Particle Image Velocimetry in PIVlab (32×32 pixel interrogation windows with 50% overlap) (55). Traction forces were computed from the displacement fields using Fourier Transform Traction Cytometry, modelling the PDMS substrate as a linear elastic, isotropic half-space. Intercellular stress tensor fields within the monolayer were subsequently reconstructed using Bayesian Inversion Stress Microscopy (BISM) (38). The isotropic (hydrostatic) stress was calculated as (σxx + σyy)/2, where positive values denote tension and negative values compression. Analysis was performed on uncropped images with free boundary condition. After, the boundary region of approximately 20% was excluded to avoid edge artifacts.

### Model

The simulations of cell colonies used an agent-based model (34, 35) with some modifications as described in *Supplementary note*. Cells were modelled as soft disks sitting on a two-dimensional substrate. Each cell was bestowed with an idealized cell cycle which is regulated by local pressure. The cell cycle in turn regulates the cell’s proliferation and growth dynamics. The cells’ positions evolve according to overdamped dynamics, since intercellular forces and substrate friction dominate inertial forces. For details, see Supplementary Information. The analysis of the simulation results was carried out in Python, making use of the Numpy and Matplotlib libraries.

## Acknowledgements

We thank Dr. Sebastian Wrighton as well as all the members of laboratory of cell and molecular mechanobiology (LCMM) at Lund University for their advice and support. Lund University Bioimaging Centre (LBIC) at Lund University is gratefully acknowledged for providing imaging resources. We also thank Willem-Jan Pannekoek for technical assistance with stencil production and cell culturing. This research was funded by the Knut and Alice Wallenberg Foundation (V.S.) via the Wallenberg Centre for Molecular Medicine, Lund; Cancerfonden (VS, 19 0445 Pj and 22 2398 Pj Projekt grant), Fru Berta Kamprad Foundation (VS, FBKS-2023-31), Crafoord Foundation (VS). S.K.S acknowledges funding from the Grants-in-Aid for Scientific Research (JSPS KAKENHI) under Grant Nos. 22K14012 and 25H01365, as well as the JSPS Core-to-Core Program “Advanced core-to-core network for the physics of self-organizing active matter” JPJSCCA20230002. Part of this research was conducted while visiting the Okinawa Institute of Science and Technology (OIST) through the Theoretical Sciences Visiting Program (TSVP) Thematic Program on Biological Information Processing (TP24BY). P.N acknowledges funding from Fru Berta Kamprads Stiftelse, and Cancerfonden. A. D. acknowledges funding from the Novo Nordisk Foundation (grant No. NNF18SA0035142 and NERD grant No. NNF21OC0068687), Villum Fonden (Grant no. 29476), and the European Union (ERC, PhysCoMeT, 101041418).

