## Supplementary figures and note for "Mechanical memory of confinement pressure governs expansion size in epithelial monolayers"

Supplementary Figure 1

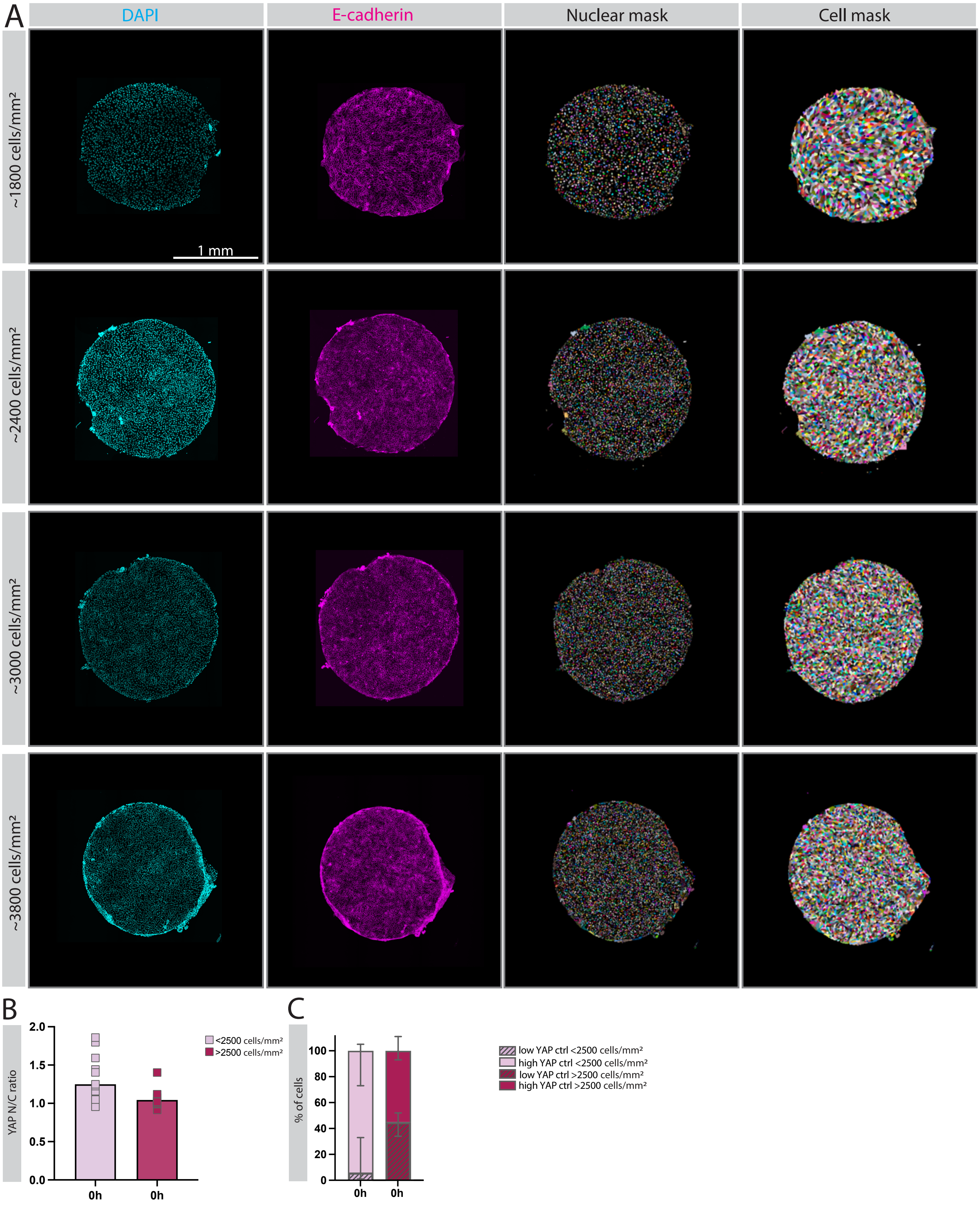

### Supplementary Figure 2

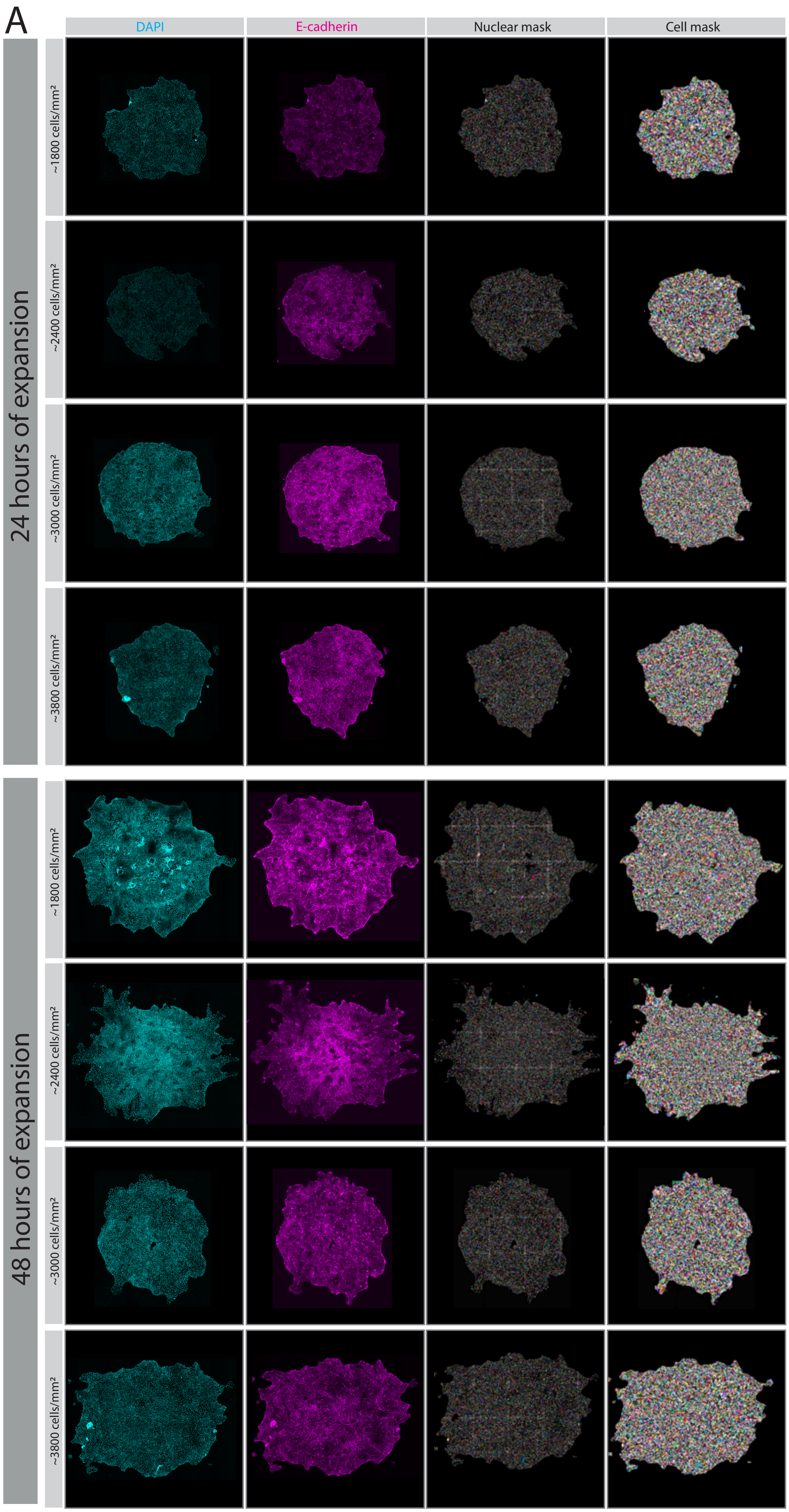

#### Supplementary Figure 3

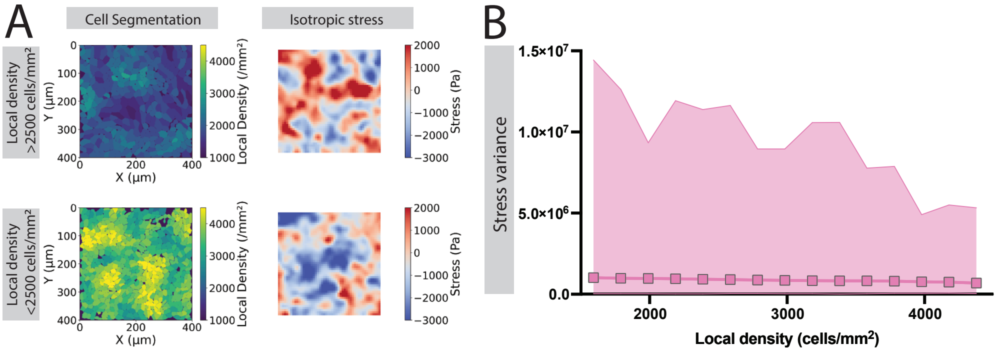

### Supplementary Figure 4

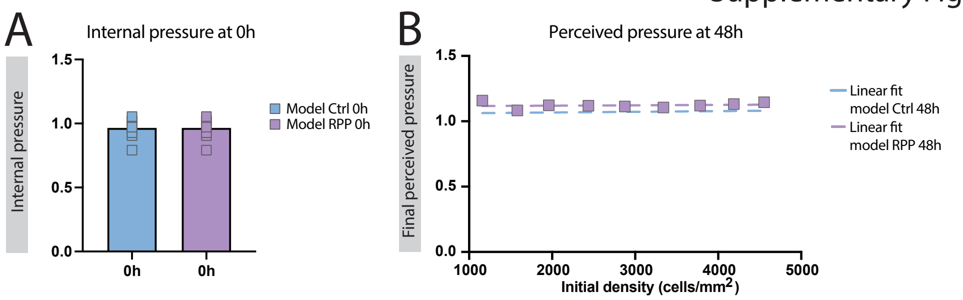

Supplementary Figure 5

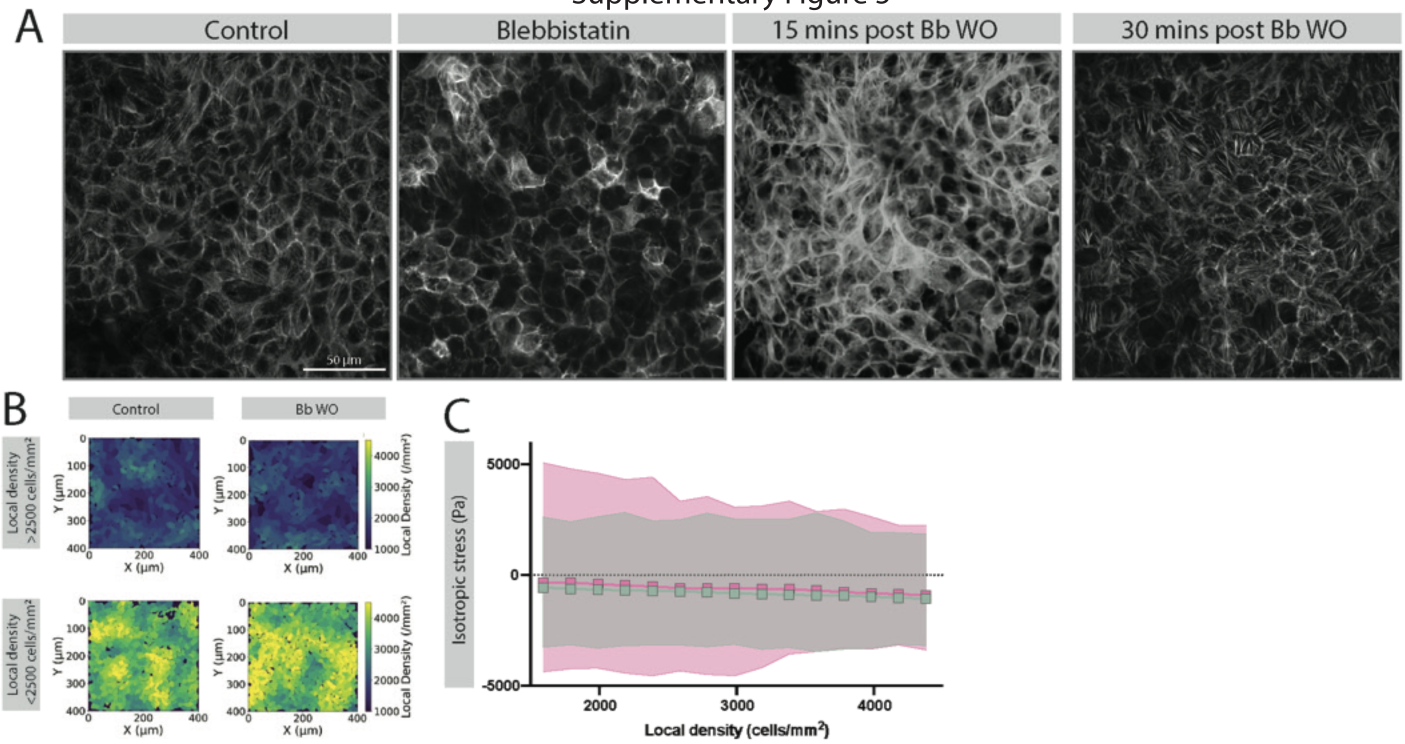

### Supplementary Figure 6

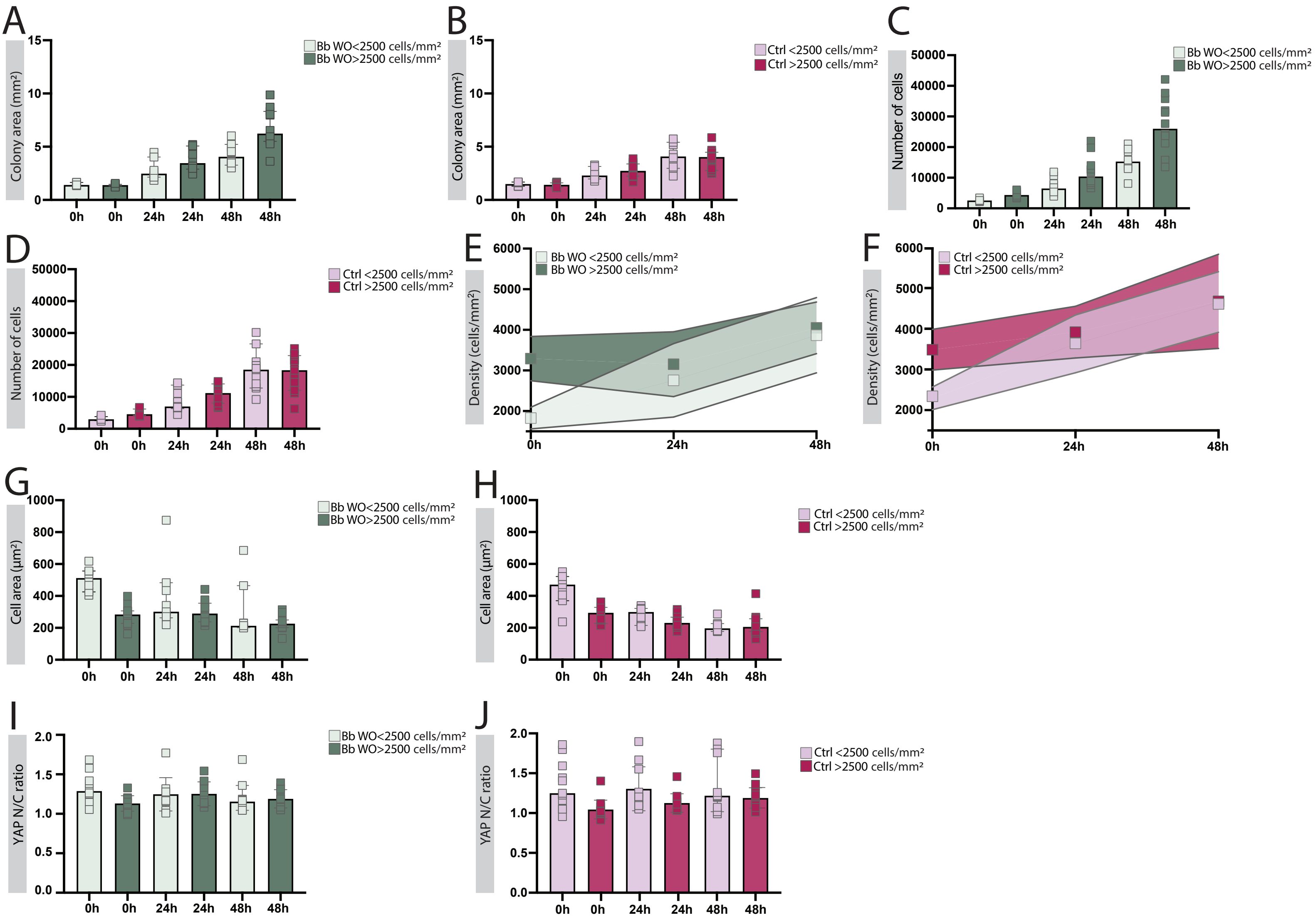

### Epithelial expansion and density homeostasis through pressure-mediated regulation of proliferation – Supplementary Information

#### 1. MODEL

The cell colony model used here is slightly modified from the model developed by Li et al. [1, 2]. Cells are modelled as soft disks sitting on a two-dimensional substrate. Each cell $i$ has a radius $R_i$ that is regulated by its cell cycle. The cell cycle determines how rapidly a cell proliferates and grows its area. Crucially, these two processes are not directly coupled but are both regulated by the cell's cell cycle activity, which in turn is regulated by pressure. In detail:

Cells interact mechanically with each other via a combination of repulsive and attractive forces. For a cell $i$ and another cell $j$, situated at positions $\vec{r}_i$ and $\vec{r}_j$, the repulsive force is assumed to be due to Hertzian contact mechanics [3],

$$\vec{F}_{ij}^{rep}(t) = \frac{f^{rep}\, h_{ij}^{3/2}(t)\, \vec{n}_{ij}}{\sqrt{1/R_i(t) + 1/R_j(t)}} \tag{1}$$

The force is proportional to the repulsive coefficient $f^{rep}$, which can in principle be calculated from the Poisson ration and elastic modulus of the cell, but is treated as a constant here, see table I. The overlap between the two cells $h_{ij}$ is calculated from the distance $r_{ij}$ between the two cells' centers

$$h_{ij} = \max(R_i + R_j - r_{ij}, 0) \tag{2}$$

with $r_{ij} = |\vec{r}_{ij}|$ and $\vec{r}_{ij} = \vec{r}_j - \vec{r}_i$. The closer cells are, the larger the overlap. When the cell-cell distance exceeds the sum of their radii, the force becomes 0. The repulsive force's direction is given by the center to center unit vector $\vec{n}_{ij} = \vec{r}_{ij}/|\vec{r}_{ij}|$. The simulation is performed for dimensionless variables, but is later matched to experimental results by Puliafito et al. [4] via the units given in table II. The attractive force $\vec{F}_{ij}^{ad}$ between cells $i$ and $j$ is modeled as an approximation of the adhesion force due to receptor-ligand interactions

$$\vec{F}_{ij}^{ad} = -f^{ad} l_{ij} \vec{n}_{ij} \tag{3}$$

The adhesive force is proportional the adhesion coefficient $f^{ad}$ and the contact length $l_{ij}$ between the cells

$$l_{ij} = \sqrt{(r_{ij}^2 - (R_i - R_j)^2)((R_i + R_j)^2 - r_{ij}^2)} \Big/ r_{ij} \tag{4}$$

The total force $\vec{F}_i$ on cell $i$ is then obtained from accounting for the interactions with all other cells

$$\vec{F}_i = \sum_{j \neq i} \vec{F}_{ij}^{rep} + \vec{F}_{ij}^{ad} \tag{5}$$

The motion of the cells is dominated by cell cell interactions and friction with the surface. As a result, we implement overdamped dynamics in which the inertia of the cells is neglected. The equation of motion for the position $\vec{r}_i$ of cell $i$ is then given by

$$\frac{d\vec{r}_i}{dt} = \frac{\vec{F}_i}{\pi R_i^2 \zeta} \tag{6}$$

We assumed here that the friction increases with the size of the cell area with the friction per unit cell area being $\zeta$. We numerically integrate eq. (6) with a time step $\Delta t$ that is small enough for numerical stability and accuracy, see table I.

Each cell $i$ contains a stylised cell cycle oscillator. The cell cycle phase $\theta_i$ is an angle that expresses the progress around the cell cycle: at cell birth, the angle is $\theta_i = 0$ and the cell divides when $\theta_i = 2\pi$. The rate at which the phase grows is given by

$$\frac{\partial \theta_i}{\partial t} = f(a_i) \tag{7}$$

with $a_i$ the cell cycle activity of the cell. We assume that $a_i \geq 0$. In the original formulation of the model, the cell cycle speed was directly proportional to the activity, $f(a) \sim a$. This meant that any decrease in activity would reduce the proliferation rate. From experimental data one can infer however, that MDCK cells exhibit a division time that is at its minimal value for a wide range of cell areas, down to $\approx 200\mu m^2$ [4]. In our model, cell division times are not determined by cell area but there is a correlation between the two quantities and which can be

Table I: Standard simulation parameters, obtained from matching to our experimental data. Units as in table II.

| Parameter | Value | Units |
|---|---|---|
| Substrate friction $\zeta$ | 0.0016 | $\tau_{\text{div}} p_r / A_q$ |
| Time step $\Delta t$) | $5 \cdot 10^{-6}$ | $\tau_{\text{div}}$ |
| Repulsive coefficient $f^{rep}$ | 17.78 | $p_r/\sqrt{A_q}$ |
| Adhesion coefficient $f^{ad}$ | 0.2 | $p_r$ |
| Cell cycle activity recovery time $\tau_a$ | 1 | $\tau_{\text{div}}$ |
| Volume recovery time $\tau_v$ | 1 | $\tau_{\text{div}}$ |
| Nominal unstressed cell area $A_1$ | 35 | $A_q$ |

Table II: Conversion between dimensionless simulation units and corresponding experimental values. Since the pressures were not matched to a particular experiment, they are kept dimensionless.

| Unit | Value |
|---|---|
| Unstressed division time $\tau_{\text{div}}$ | 18h |
| Quiescent cell area $A_q$ | 30 µm$^2$ |
| Reference pressure $p_r$ | 5 |

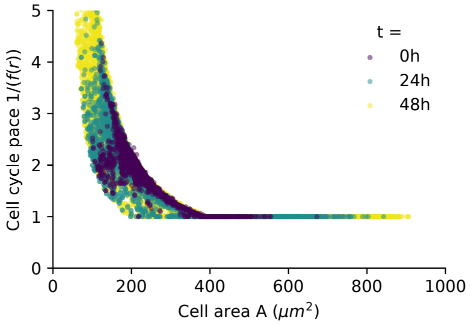


Figure 1: **Cell cycle pace vs. cell area.** The cell cycle pace $1/f(a)$ is an estimator of the expected cell division time (which can exceed the duration of the simulation). A value of $1/f(a) = 1$ corresponds to a division time of $\tau_{\text{div}}$.

directly compared to the experimental data. The original model already yielded an increase in division time for cells with areas well above $200\mu m^2$. We address this qualitative difference by adjusting the regulation function such that the division time is minimal as long as the activity $a$ is above a saturation activity $a_{sat}$,

$$f(a_i) = \frac{2\pi}{\tau_{\text{div}\,i}} \min\left(\frac{a_i}{a_{sat}}, 1\right) \tag{8}$$

with saturation activity $a_{sat} = 1/3$. This yields qualitatively the correct pace of divisions, see fig. 1. The division time $\tau_{\text{div}\,i}$ of an unstressed cell ($a = 1$) is drawn from a normal distribution with mean $\tau_{\text{div}}$ and standard deviation $\tau_{\text{div}}/3$.

The cell cycle is assumed to be regulated by pressure, the isotropic contribution to the stress tensor. We estimate the pressure acting on cell $i$ in the two-dimensional plane of the cell colony as the sum of all the inward-pointing cell forces divided by the cell's circumference

$$p_i = \sum_{j\neq i} \frac{-(\vec{F}_{ij}^{rep} + \vec{F}_{ij}^{ad}) \cdot \vec{n}_{ij}}{2\pi R_i(t)} \tag{9}$$

The pressure then regulates the activity via

$$\frac{1}{a_i}\frac{\partial a_i}{\partial t} = \frac{1}{\tau_a}\left(1 - a_i - \frac{p_i}{p_r}\right) \tag{10}$$

The cell cycle activity reacts to mechanical pressure on a time scale of $\tau_a$, whereas the reference pressure $p_r$ sets the scale of pressures that will lead to quiescence ($r \to 0$) over long times.

In between divisions, the area of a cell is regulated by cell activity as

$$\frac{1}{V_i}\frac{\partial V_i}{\partial t} = c\, a_i - \frac{1}{\tau_v}\frac{V_i - A_q}{A_q} \tag{11}$$

The first term describes cellular growth as proportional to a growth rate $c$ and the cell activity. From an analysis of the stationary state of an unstressed cell we connect the growth coefficient $c$ with the average cell area of an unstressed cell $A_1$, via $c = \ln(2) + (A_1 - A_q)/(\tau_v A_q)$. The second term describes the tendency of cells to reach a small quiescent volume if the cell cycle arrests, which occurs here on a time scale of $\tau_v$. When a cell divides, it is replaced by two daughter cells. Each daughter cell obtains half the area of the mother cell. Both daughters are positioned at the maximum possible distance along a random axis within the mother's footprint. Their cell cycle activities are inherited from the mother, their cell cycle phases are reset to 0. The unstressed cell division times $\tau_{\text{div}\,i}$ are randomly drawn from the distribution given above.

#### 2. MATCHING THE SIMULATIONS TO PULIAFITO ET AL (2012)

The model and parameter values as described in [1] only provided a qualitative match to the MDCK experiments by Puliafito et al. [4] when growing a colony from a single cell. However, a near quantitative match is achieved by increasing the maximum possible size of cells, see table I and table II, reducing the friction coefficient to $\zeta = 0.0004$ as well as using the adjusted cell cycle regulation function eq. (8). Note that we used a smaller value for the friction coefficient than the one mentioned in table I (which is the one used for matching to our own experiments), as is discussed in the next section.

#### 3. MATCHING THE SIMULATIONS TO OUR EXPERIMENTS

First, we simulate stenciled colonies initialised from a single cell in the center of a circular region of radius $R_{wall}$. The confinement is achieved with a Lennard-Jones type force along the region's perimeter. We assume that the confinement is centered on the origin of the simulation box. For a cell $i$ at position $\vec{r}_i$ within the confined region, the confinement wall is at a distance $d(\vec{r}_i) = R_{wall} - |\vec{r}_i|$ and exerts a force

$$\vec{F}_{wall}(\vec{r}_i) = \frac{-\epsilon}{4R_i^2}\left(2\left(\frac{2R_i}{d(\vec{r}_i)}\right)^{14} - \left(\frac{2R_i}{d(\vec{r}_i)}\right)^{8}\right)\frac{\vec{r}_i}{|r_i|} \tag{12}$$

if $d < r_{cut} = 2^{7/6}R_i$ and 0 else. The cutoff $r_{cut}$ is selected such that the wall is purely repulsive. Confinement parameters are $R_{wall} = 91\sqrt{A_q}$ and $\epsilon = 5$. The simulated colony at first expands outward until it reaches the confinement wall and then becomes denser and less active over time, see fig. 3.

Then, we extract snapshots from that simulation at appropriate densities to be used as initial configurations for new simulations in which the circular confinement is

Firstly, for a snapshot of the stenciled simulation, we fixed the cells in place, froze their cell cycle phase $\theta_i$, turned off division and growth mechanisms. The cell cycle was then assumed to react to a reduced perceived pressure instead of the actual mechanical pressure. We assumed a reduction in the perceived pressure of each cell $p_{red,i} = p_i \Delta_p$ with $\Delta_p = 0.6$ for a duration of $\Delta_t = \tau_{\text{div}}/6$ (equivalent to 3h of experimental time). With these perceived pressures, we updated the cell cycle activities $a_i$ by numerically integrating eq. (10) forward in time for a duration of $\Delta_t$. This yielded the modified activities denoted as RPP in Fig. 4B.

Secondly, we used the snapshots with these updated activities as initial states for simulations without confinement. Assuming that the perceived pressures would not immediately match the actual pressures, but instead exponentially decay towards them, we made the cell cycle be regulated by the dynamically scaled perceived pressure as follows,

$$p_{red,i} = p_i \left(1 - (1 - \Delta_p) e^{-t/\tau_p}\right) \tag{13}$$

with $\tau_p = \tau_{\text{div}}$.

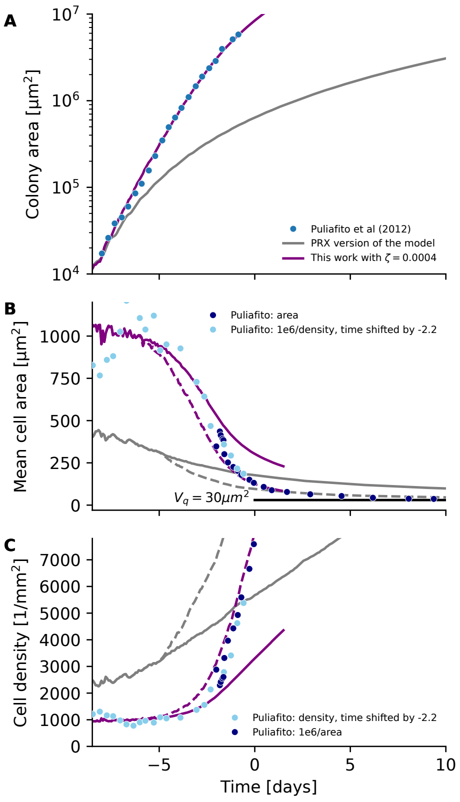


Figure 2: **Matching our simulations to experiments by Puliafito et al. [4].** (A) Total area of the colony, (B) mean cell area, and (C) cell average cell density for experimental and simulation data over time. The experimental data by Puliafito et al. [4] was extracted by hand using Webplotdigitizer [5]: Colony areas were extracted from their Fig 1B, cell density from Fig 1C and the median of the cell size distribution (called cell area here) from Fig 3C. Their density data is from the inner region of the colony. Their cell area data is from an unspecified region of the colony. We found that density and median cell area could be made consistent when shifting the density data by -2.2 days. The simulations were grown from a single cell. The model with parameters as reported in [1] (labelled PRX version) qualitatively but not quantitatively reproduces this experimental data. In order to achieve quantitative fit, we adjusted the simulation parameters, see table I. The mean cell area and cell density are shown for as colony averages (solid lines) and for the center of the colony (dashed lines). The latter is more comparable to the approach chosen by Puliafito et al. [4] and thus yields a more fitting comparison.

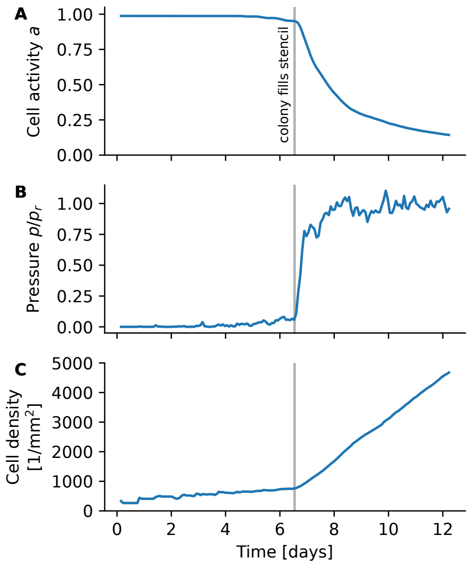


Figure 3: **A colony seeded from a single cell expands to fill a circular stencil.** (A) Cell activity $a$, (B) average pressure $p$ (normalised by the reference pressure $p_r$) across the colony, and (C) average cell density as a function of time are plotted for simulation data of a single cell being seeded in the center of a circular stencil confinement. The grey vertical line indicates when the colony first fills the stencil completely. Once the colony fully fills the stencil, pressure rises rapidly, leading to reduced cell activity over time. While the colony area cannot exceed the available stenciled area, the cells keep dividing and thus increase the density of the colony over time.

removed. This is in analogy to removing the stencil in the experiments. These simulations are then monitored for the equivalent of 48h of experiment time.

We find that our experimental systems in general expand less quickly than the colonies by Puliafito et al. [4] when compared at similar colony areas. We note that our systems have higher density at the same colony area, but since we demonstrate in this work that cell expansion is largely density independent, this should not play a role. One reason could be that the used substrate is different. In our simulations, this can is addressed by increasing the friction coefficient with respect to the value used for matching to the Puliafito data. Find our typical parameter values in table I.

##### 3.1. Reduced Perceived Pressure (RPP)

The reduction of the pressure as perceived by the cell cycle was modelled in two ways.
